# Interpretable Machine Learning for Perturbation Biology

**DOI:** 10.1101/746842

**Authors:** Bo Yuan, Ciyue Shen, Augustin Luna, Anil Korkut, Debora S. Marks, John Ingraham, Chris Sander

**Author notes:** Joint first authors (Bo Yuan, Ciyue Shen). Correspondence to reaches the principal authors.

## Abstract

Systematic perturbation of cells followed by comprehensive measurements of molecular and phenotypic responses provides an informative data resource for constructing computational models of cell biology. Models that generalize well beyond training data can be used to identify combinatorial perturbations of potential therapeutic interest. Major challenges for machine learning on large biological datasets are to find global optima in an enormously complex multi-dimensional solution space and to mechanistically interpret the solutions. To address these challenges, we introduce a hybrid approach that combines explicit mathematical models of dynamic cell biological processes with a machine learning framework, implemented in Tensorflow. We tested the modeling framework on a perturbation-response dataset for a melanoma cell line after drug treatments. The models can be efficiently trained to accurately describe cellular behavior, as tested by cross-validation. Even though completely data-driven and independent of prior knowledge, the resulting *de novo* network models recapitulate some known interactions. The main predictive application is the identification of combinatorial candidates for cancer therapy. The approach is readily applicable to a wide range of kinetic models of cell biology.

## Introduction

The emergence of resistance to single anticancer agents has highlighted the importance of developing combinations of agents as a more robust therapeutic approach to cancer treatment (Fitzgerald *et al*., 2006; Garraway and Jänne, 2012; Mansoori *et al*., 2017; Mokhtari *et al*., 2017). However, experimental screening of all possible pairwise or higher-order combinations of currently available agents is practically unrealistic. The space of potential new therapeutic targets is even larger and more challenging to explore experimentally. To efficiently narrow down the search space and nominate promising sets of experimentally testable candidates, computational models have been used to predict cellular responses based on sets of perturbation experiments (Azmi *et al*., 2010; Ryall and Tan, 2015; Cheng, Kovács and Barabási, 2019), but these have been limited in scope. The ability to model cell biology at a larger scale and to infer causal mechanisms to generalize to unobserved perturbations is critical in facilitating the search for combinatorial, potentially therapeutic candidates.

### Perturbation-response profiling in cell biology

In order to understand cell behavior, a large variety of experimental approaches has been used to profile cellular responses under different perturbations. Biochemical and cell biological experiments testing relationships of particular protein-protein pairs have, for many years, been successfully used to identify signaling cascades (Cerami *et al*., 2011; Wrzodek *et al*., 2013; Croft *et al*., 2014), but one-by-one perturbation experiments are laborious and the resulting models, while insightfully descriptive, typically do not provide the ability to quantitatively predicting detailed molecular or system-level responses. Phenotypic screening collects high-throughput information on whole-cell responses with univariate readouts such as cell viability or growth rate (Lamb *et al*., 2006; Cheung *et al*., 2011; Wang *et al*., 2014; McDonald *et al*., 2017; Tsherniak *et al*., 2017). In order to resolve intracellular interactions and provide mechanistic insights, systematic methods have been developed to profile post-perturbational molecular responses with increasing coverage, e.g., changes in transcript (Dixit *et al*., 2016; Niepel *et al*., 2017; Norman *et al*., 2019) and protein (Korkut *et al*., 2015; Hill *et al*., 2017) levels. These rich datasets challenge computational methods to describe mechanisms more comprehensively and to quantitatively model cell responses to unseen perturbations, e.g., for the design of experiments to test mechanistic hypotheses or for the design of combinatorial therapeutic interventions.

### Computational modeling

Various computational methods have been developed to infer interactions and predict cellular responses (Vanhaelen, Aliper and Zhavoronkov, 2017). Static models use, e.g., co-expression models (Carter *et al*., 2004; Wang *et al*., 2009; Babur *et al*., 2010), maximum entropy networks (Lezon *et al*., 2006; Locasale and Wolf-Yadlin, 2009), or mutual information related methods (Meyer, Lafitte and Bontempi, 2008; Chan, Stumpf and Babtie, 2017), to construct network models of molecular interactions (Şenbabaoğlu *et al*., 2016; Yi *et al*., 2017), or use regression models to directly predict cellular responses based on molecular perturbation-response measurements (Dixit *et al*., 2016; Norman *et al*., 2019). On the other hand, dynamic models, such as Boolean network models (D’haeseleer, Liang and Somogyi, 2000), fuzzy logic models (Aldridge *et al*., 2009), dynamic Bayesian networks (Zou and Conzen, 2005) and ordinary differential equation (ODE) network models (Nyman *et al*., 2019; Gardner *et al*., 2003), can provide mechanistic insight in terms of propagation of cellular signals to phenotypic response over time, but typically require prior knowledge of interaction parameters and thus currently only work for small systems (Gardner *et al*., 2003; Klinger *et al*., 2013). New algorithms to parametrize large-scale mechanistic models, while computationally efficient, require prior knowledge of a set of relevant interactions (Fröhlich *et al*., 2018). Insufficient prior knowledge is one of the major constraints for modeling large systems, e.g., in that prior information is not available for all components or is aggregated from disparate experimental sources and thus lacks uniform context. A more rigorous approach is to use uniform datasets generated in systematic experiments in one experimental context and then perform de novo structure inference of an interaction network specific for that context. Given such data for large systems, the computational challenge is to search for optimal interaction parameter sets in a complex multi-dimensional solution space. Previous dynamic optimization approaches such as Monte Carlo (MC) methods and belief propagation (BP) algorithms, have been used to construct data-driven network models (Bruggeman *et al*., 2002; Nelander *et al*., 2008; Hug *et al*., 2013; Klinger *et al*., 2013; Korkut *et al*., 2015). Still, these may not efficiently scale to larger systems (e.g., MC) or may require excessive approximations for the chosen mathematical model to facilitate efficient exploration of solution space (e.g., independent row approximation in BP) (Bruggeman *et al*., 2002; Korkut *et al*., 2015). Therefore, to achieve good accuracy of parameter inference for larger systems and to gain the ability to generalize to more sophisticated kinetic models, a more general and potentially more powerful data-driven modeling framework is needed.

### Machine learning and interpretability

Recently, deep learning has become an effective data-driven framework capable of generating predictions for large and complex systems. Gradient descent implemented with automatic differentiation, which has been broadly used in training graphical models, allows efficient parameter optimization in complex network systems. This framework has been successfully applied to many domains of biomedical research, from pathology image classification (Hou *et al*., 2016; Esteva *et al*., 2017) to sequence motif detection (Zhou and Troyanskaya, 2015). While predictive power of deep learning models is often impressive, their interpretation, which is crucial for providing understandable and, therefore, more trustable predictions, remains challenging. The complex multi-layer network architecture of most deep learning models lacks explicit representations and consequent direct interpretation. This difficulty is sometimes called the “black box” problem (Montavon, Samek and Müller, 2018). To address this problem, we apply a deep learning optimization approach to learn a data-driven model (called “CellBox”) that incorporates an explicitly interpretable network of interactions between cellular components, instead of a black-box neural network, while aiming to maintain a high level of learning performance.

CellBox is designed to be a framework for computational modeling of cellular responses to perturbations that i) links perturbations to molecular and phenotypic changes in a unified computational model; ii) quantifies time-dependent (dynamic) cellular responses; iii) promises training efficiency and scalability for large-scale systems; iv) is interpretable in terms of interactions that can be compared to established models of molecular biology, such as signaling pathways. Here, we construct a non-linear ordinary differential equation (ODE) based model representing a biological network of 99 components connecting perturbations, protein response, and phenotypes to simulate dynamic cellular behavior. The network connections are directly learned from post-perturbational data under 89 experimental conditions with the objective of accurately reproducing the molecular and cellular responses on training data. To reach this objective, we implemented gradient descent with automatic differentiation to infer interaction parameters in the ODE network, which can then be exposed to novel perturbations to predict cell behavior. The key performance criterion for the data-driven model trained with a relatively small set of experiments is whether the model is able to provide reasonably accurate predictions on a large set of unseen perturbation conditions. Anticipating the availability of increasingly informative perturbation-response data sets in diverse areas of cell biology, we present CellBox as a generally applicable framework for modeling a broad range of dynamic cell behavior.

## Results

### CellBox model of perturbation biology

In order to construct a data-driven model to predict the dynamics of molecular and cellular behavior under combinations of drug treatments, the perturbation data has to have 1) paired measurements of changes in protein levels and cellular behavior for a set of perturbations; and 2) training and withheld data to test model performance. Here, we used a perturbation dataset for the melanoma cell line SK-Mel-133 (Korkut *et al*., 2015), which contains molecular and phenotypic response profiles of cells treated with 12 different drugs and their pairwise combinations (Figure 1a, S1). For each of the 89 perturbation conditions, levels of 82 selected proteins and phosphoproteins were measured in cell lysates before and 24 hours after perturbation on antibody-based Reverse Phase Protein Arrays (RPPA). In parallel, cellular phenotypes were assayed, including cell cycle progression and cell viability. With parallel measurements of proteomic and phenotypic responses to a systematic set of perturbations, this dataset provides sufficient information to construct network models that quantitatively link molecular changes to cellular responses.

**Figure 1.**
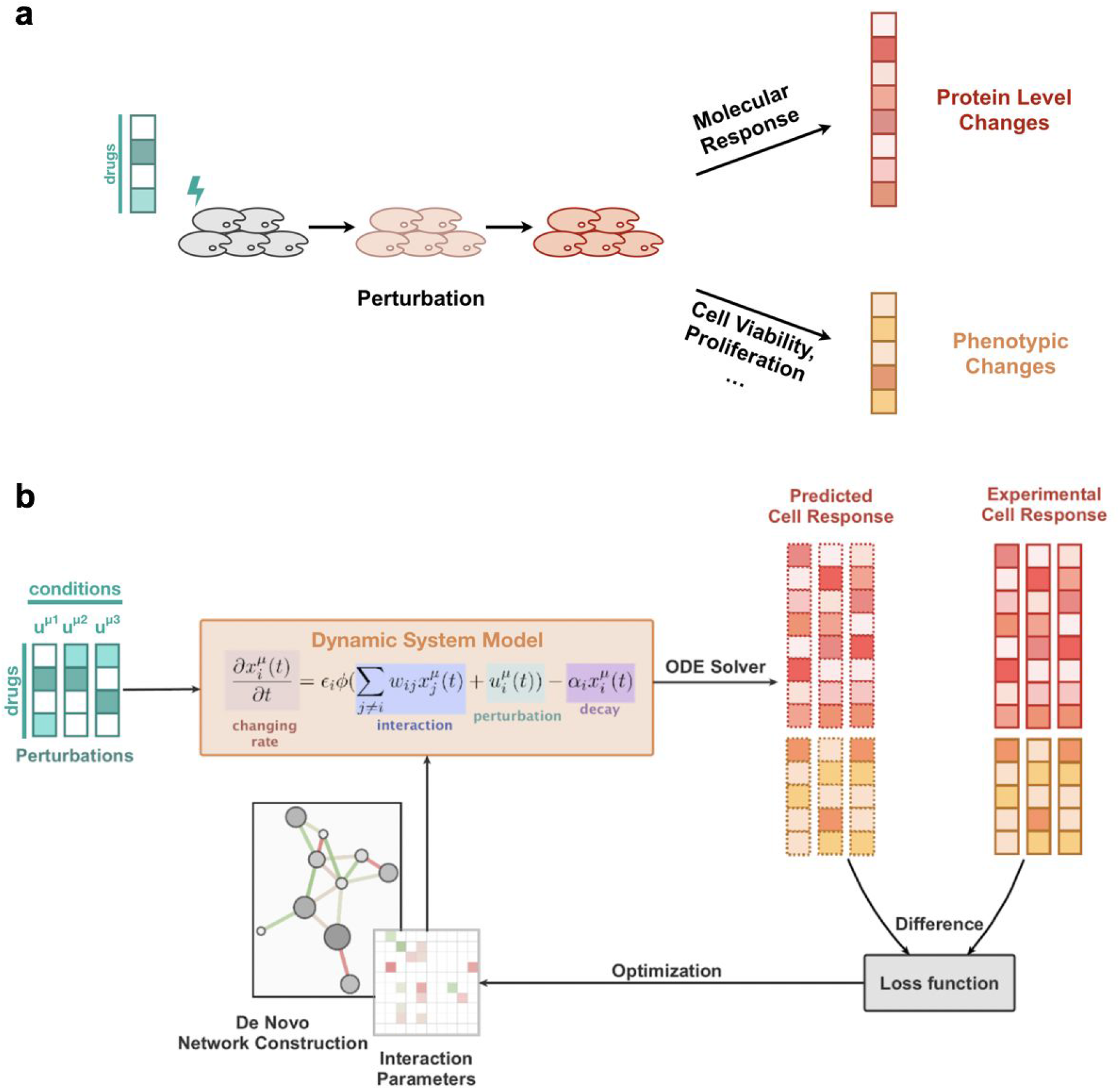
CellBox: dynamic modeling of cellular systems with perturbation data. **a.** Perturbations such as drugs are used to disturb the cellular system. The cell responses, including protein and phosphoprotein level changes, and phenotypic changes, were measured to provide information for model construction. **b.** Systematic responses of the cells under various drug perturbations were used to construct an interpretable machine learning model. CellBox models system behavior in terms of interaction parameters connecting molecular (proteins and phosphoproteins) and phenotypic variables using a set of differential equations. CellBox was trained iteratively by optimizing interaction parameters to fit the numerically simulated system response to experimental observations. After training on pairwise data of input perturbation and output system behaviors, the CellBox model can be used to predict the cellular response to arbitrary perturbation conditions.

We used a set of ordinary differential equations (ODEs with a non-linear envelope) (Figure 1b) to model the dynamic responses of the system to drug perturbations (See Methods). The parameters of the ODEs (*w_ij_*, ∼10,000 in total) are the interaction strengths between the entities in the network model. The simplicity of the interaction dynamics (Figure 1b), the non-linear envelope, as well as the restoration term − α*_i_ x_i_* (*t*) are computational devices, roughly analogous to mean-field approaches, to account for the fact that the data is limited to a relatively small fraction of all cellular components and to avoid instabilities (Nelander *et al*., 2008; Molinelli *et al*., 2013; Korkut *et al*., 2015). The interaction parameters were randomly initialized and updated throughout the model training process, with the objective to minimize a loss function. For the loss function, we chose the Euclidean distance between experimental data and the results of the numerical simulation of the ODE model, plus an L1 regularization penalty on network density to avoid overfitting (See Methods, eqn. 3). We used Heun’s ODE solver (Süli and Mayers, 2003) to numerically simulate the ODE system and the Adam optimizer (Kingma and Ba, 2014) with automatic differentiation to minimize the loss function. Taken together, we constructed an ODE model of a cell biological system trained using perturbation data, which we named CellBox.

### CellBox can be trained on perturbation data to accurately predict cell response

In order to test the prediction performance of this training scheme, we randomly selected 70% of the perturbation data (n = 62 conditions) for training and withheld the rest 30% (n = 27 conditions) for testing. 20% of the training data was used as a validation set to stop model training when the performance on the validation set did not further improve. We manually fine-tuned the hyperparameters, including learning rate, regularization, and ODE simulation time, to increase the training efficiency (Figure S2). At the end of the training, the numerical solutions of the ODE model converged efficiently to experimental data (Figure 2a, 2b). We repeated the modeling scheme with 1,000 independent random data partitions to construct models for each partition. The average predictions on test sets across all models and all conditions correlate with experimental data with a Pearson’s correlation coefficient of 0.94 (Figure 2c). A more refined analysis of individual perturbation conditions showed that the model trains equally well for all conditions (Figure 2d), and that model performance is independent of data scaling normally applied to protein profiling data (Figure S3). The results illustrate that the CellBox model can be efficiently trained with perturbation data to predict cell response to experimentally applied perturbations accurately.

**Figure 2.**
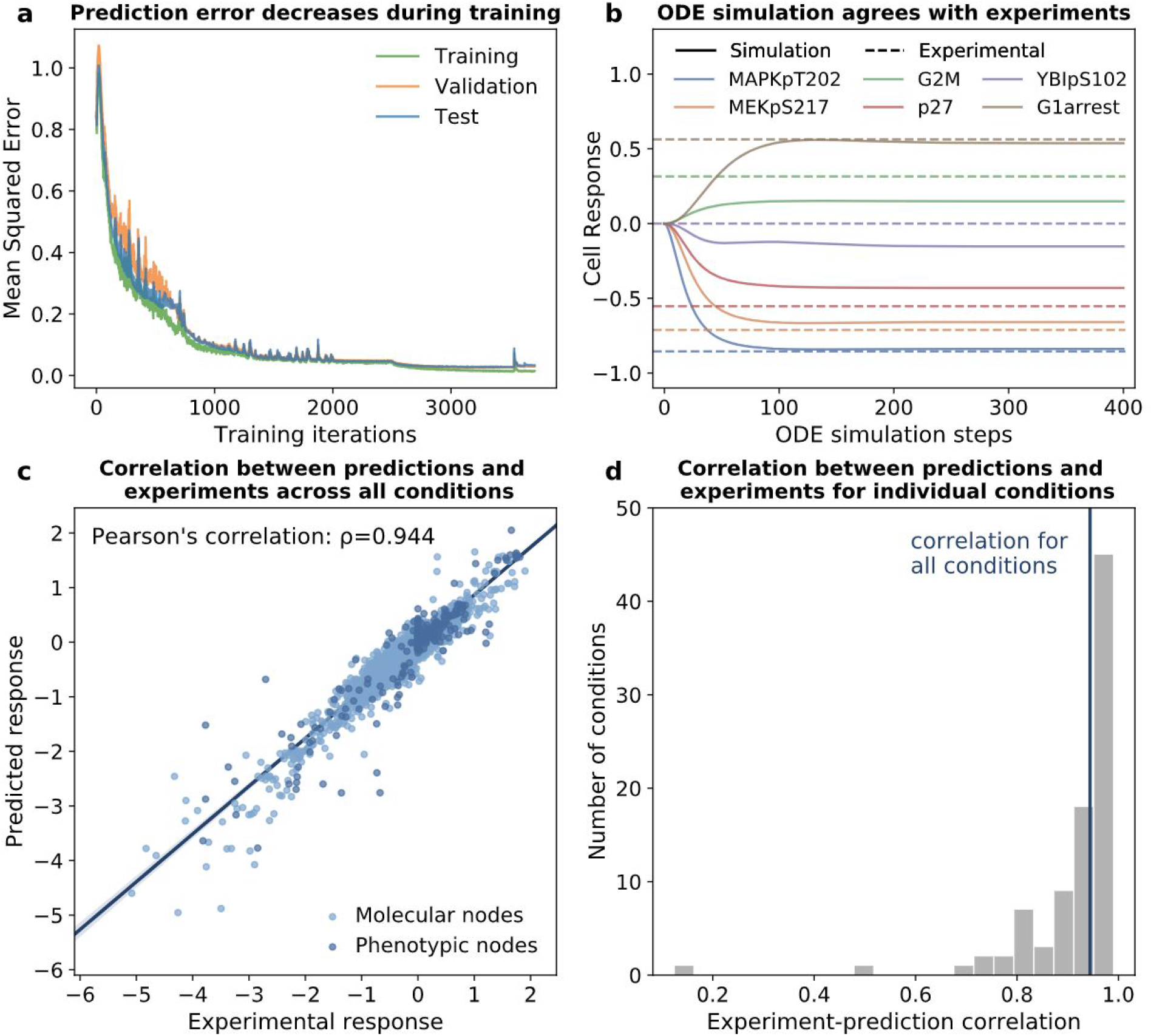
CellBox convergence and prediction accuracy on randomly partitioned training-test datasets. **a.** Over training iterations, the mean squared error on the training set (56% of the entire dataset), validation set (14%) and test set (30%) decreased nearly monotonically, and the models converged at the end of the training. **b.** The predicted molecular and phenotypic responses at the steady state of the ODE simulations agree with the experimental data on the test set. A subset of molecular measurements (MAPKpT202, YB1pS102, MEKpS217, and p27) and phenotypic measurements (G2M and G1arrest) are shown. Cell response is defined as the log_2_ ratio of post- and pre-perturbation measurements. The annotations and the full set of measurements are in Supplementary Table S1. **c.** Across 1,000 models trained with different data partitions, the average predicted responses correlate with experimental observations (Pearson’s correlation ρ = 0.944, regression line in dark blue with 95% confidence interval). Each point represents one measurement, either molecular or phenotypic, in one perturbation condition. **d.** Nearly all predictions for individual conditions have high correlations with experimental measurements.

Even though ∼70% of the models reached steady solutions of the ODEs (Figure 2b), some models converged to oscillatory solutions (Figure S4a). In order to test whether the oscillation is an artifact of data partitioning during model training, we re-trained the models with the same train-test data partitioning but multiple different random seeds for the computational optimizer (See Methods). For each partition of the training data, both steady and oscillatory solutions can result (Figure S4a-d), and the emergence of oscillatory solutions is independent of ODE solvers (Figure S4e-f). Based on the assumption that the population average of cell response reaches a stable and non-oscillating steady state 24 hours after drug treatment, we excluded the oscillatory models in the following analysis (See Methods). Taken together, these results indicate that CellBox, a data-driven ODE-based cellular system model, can be trained to accurately predict dynamics of cell response, without any requirement of prior knowledge about the relationship between particular protein levels or phenotypes.

### CellBox model predicts cell response for single-to-combo and leave-one-drug-out cross-validations

Even though the model makes accurate predictions with different training data, data partitioning, especially random partitioning, raises the concern of information sharing between training and test datasets. Combinatorial conditions in both datasets might share the same drugs such that the test set might not be truly independent of training and, therefore, is suboptimal for rigorous evaluation of the model performance. Moreover, the ability to predict the combinatorial effect of a drug, e.g., dominant, additive, synergistic, when none of its combinations has been seen by the model, is a non-trivial challenge in the context of making accurate predictions of experimentally untested drug combinations.

In order to address these points, rather than training the model with random data partitioning, we instead designed more rigorous tasks: single-to-combo (Figure 3a, S7) and leave-one-drug-out cross-validation (Figure 3b, 3c) for each drug. In the single-to-combo analysis, all single-drug treatment conditions were used for training, and predictions were made on all combinatorial drug conditions. In leave-one-drug-out cross-validation, all the combination conditions containing the treatment of a particular drug with or without the corresponding single drug conditions were withheld while the rest of the conditions were used for training. In these more stringent tests, we found that the predicted values for withheld data were still highly correlated with the experimental observations (average Pearson’s correlation: 0.93 for single-to-combo; 0.94 for leave-one-drug-out with single conditions, similar to that of the training with random partition; 0.79 for complete leave-one-drug-out). Under all three scenarios, on this dataset, CellBox outperforms the belief propagation (BP) dynamic model approach previously used in perturbation biology (Korkut *et al*., 2015) in terms of predictive accuracy. These results indicate that the CellBox model can be trained with a relatively small set of perturbation data and that its predictions can be generalized to unseen combinatorial perturbations. In particular, CellBox models predict more accurately than linear models in the single-to-combo scenario (Figure S7), suggesting that CellBox can capture the non-additive (synergistic or antagonistic) effects, which is particularly useful in nominating therapeutic drug combinations.

**Figure 3.**
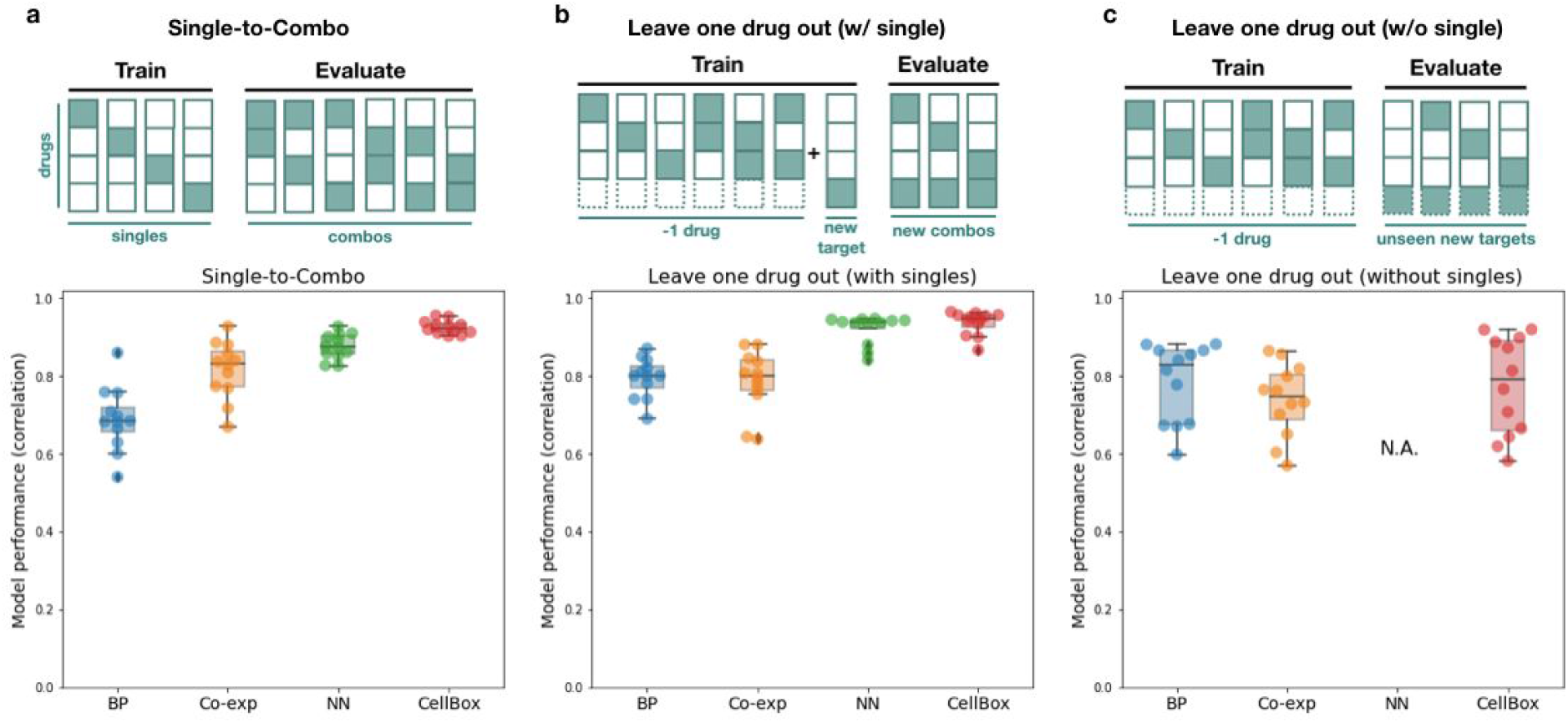
CellBox can accurately predict cell response for single-to-combo and leave-one-drug-out cross-validations. **a.** When only single conditions were used for training (single-to-combo), the CellBox models predict the effects of combinatorial conditions with high accuracy and outperform the dynamic network model inferred using belief propagation (BP), the static co-expression network model (Co-exp), and a neural network regression model (NN) trained on the same data. **b.** When combinatorial conditions associated with one drug were withheld from training, the CellBox models retain high accuracy for predicting the effects of unseen drug pairs. **c.** When all conditions associated with one drug were withheld from training, the ODE network models predict the effects of the withheld drug with reduced accuracy, but direct regression models such as NN cannot generalize to unseen targets at all. For each model type, the performance was evaluated by Pearson’s correlation between predicted cell response and experimental cell response.

CellBox models are dynamic network models of a cell biological system. To test whether such interpretable network models of molecular interactions help increase model predictive power, we compared the results to those of a static biological network model and a deep neural network model. The static network model was constructed by learning co-expression correlation for each pair of protein nodes (Co-exp) while the deep neural network model was trained to directly regress phenotypic changes against parameterized perturbations (NN) (see Methods). In all three tasks, the static network models had lower accuracy relative to the dynamic CellBox. The NN had comparable performance to CellBox in the cross-validation for individual drugs, but its performance dropped significantly in the single-to-combo analysis (Figure 3a). Furthermore, the NN was unable to generalize to unseen targets whose information is completely excluded from training (Figure 3c, Figure S6). Taken together, due to the lack of mechanistic and dynamic information, static network or direct regression models appear to be less suitable for facilitating the search for combinatorial targets.

### Model performance is robust against noise and reduced training set size

To examine model robustness of the CellBox models against a reduction in training data, we tested the stability of model performance when the data quality or quantity is compromised. To test the former, we introduced different levels of multiplicative Gaussian noise (see Methods) into the input molecular and cellular response data and trained models on the resultant noisy datasets. The assumption behind such multiplicative noise is that the uncertainty and noise in experimental measurements arise around the true values. When comparing the predicted response in test sets to the experimental data, we found that the predictions from training on the noisy data retain similarly high correlations to experimental data as those trained on the original data, even with the addition of 5% multiplicative Gaussian noise (Figure 4a). As the magnitude of the noise increases, the model performance decreases gradually in terms of both convergence (Figure S8a) and predictive power (Figure 4a). We observed similar behavior when challenging the CellBox models with additive Gaussian noise (Figure 4b, S8b). The predictivity of CellBox models can tolerate an addition of up to σ*_add_* = 0.20 additive Gaussian noise, i.e., about half the data standard deviation σ*_data_* = 0.46. We conclude that model performance is stable in the presence of moderate experimental errors.

**Figure 4.**
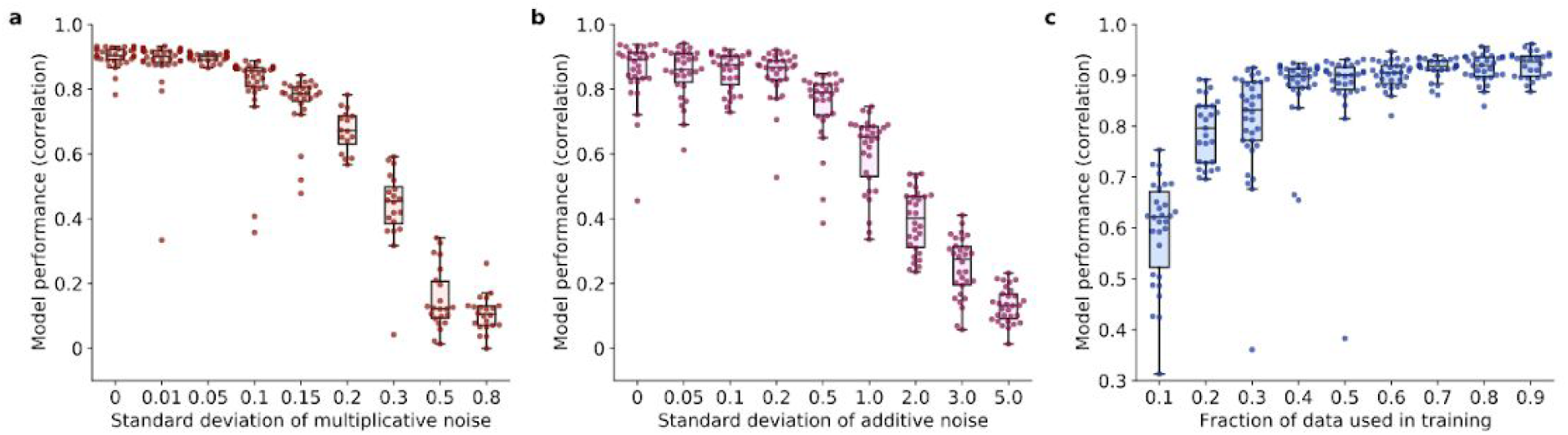
Model performance is stable against data noise and data reduction. **a-b.** Correlation between predicted responses and experimental responses in the test set decreases as an increased level of multiplicative noise (**a**) or additive noise (**b**) is added to the training data (each dot represents one model). The CellBox models can tolerate up to σ*_mul_* =0.05 multiplicative noise or σ*_add_* =0.20 additive noise. **c.** Correlation between predicted responses and experimental responses in the test set increases with an increasing quantity of data used for model training. For the current dataset, the correlation plateaus when 40% of the original dataset is used.

To test the dependency of model performance on data quantity, we trained the model on subsamples of the experimental dataset. We trained models with varying amounts of data (from 10% to 90% in steps of 10%) and found that the models could make accurate predictions of withheld data with as little as 40% of the complete dataset (Figure 4c). We found that increasing the size of the training set further has diminishing returns in terms of model performance. This implies, on the dataset used here with an interaction network of ∼100 components, that a comparatively small number of perturbation conditions (40-100, rather than directly testing all ∼3,000 possible combinations) are sufficient for constructing reasonably predictive models. This example may be a useful guide for power calculations for systems with hundreds of measured components, which would be of considerable interest.

### Comparison of the network models with prior knowledge about pathways

We used ordinary differential equations as the core framework of the current version of the CellBox mathematical model. Each parameter in the model represents the strength and direction of a biological interaction. In order to investigate whether the inferred interactions are consistent with current knowledge of biological pathways, we used the entire dataset as training data to generate 1,000 full models and examined the resulting *de novo* network edges learned from training. In order to measure both the strength and stability of edge inference, we used t-scores (see Methods) to assess the statistical significance of each interaction, where a higher absolute t-score indicates higher interaction strength and lower variance across the models (Figure 5a). Using the primary targets of the drugs as the ground truth, we first examined the interactions between the drug-activity nodes and their downstream effectors. We found that all 12 drug-activity nodes had significant edge connections to their known primary downstream protein effectors with the interaction directions consistent with their expected effects (Figure 5b), suggesting the models were able to capture the literature-provided interactions between the drug target and their downstream effectors.

**Figure 5.**
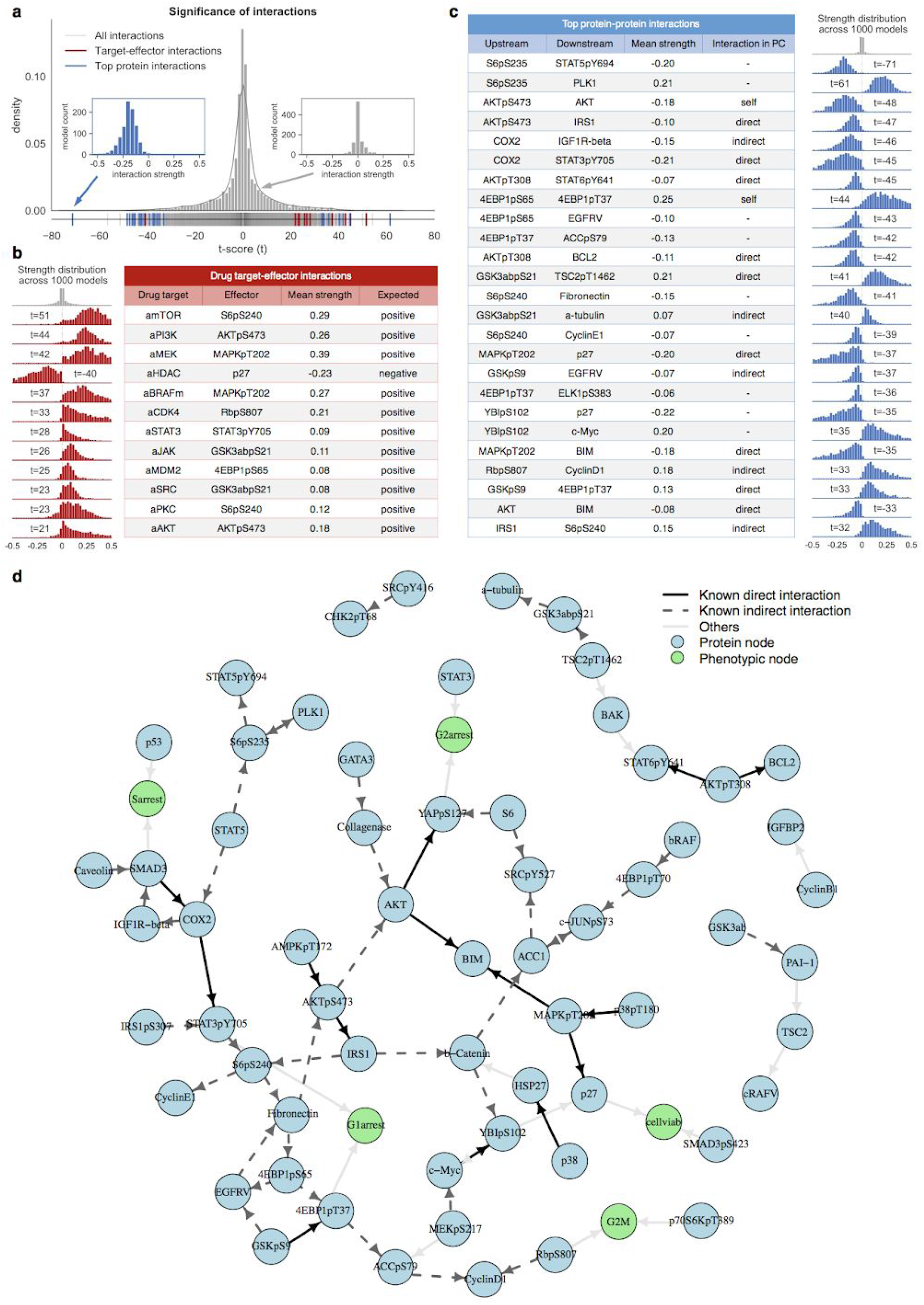
Comparison of the network models with prior knowledge about pathways. **a.** The t-score distribution of all interactions across 1,000 full models suggests that a small fraction of interaction strengths is significantly different from zero. Insets are two examples of interaction strength distributions across models. **b.** All 12 interactions between drug target (drug activity nodes) and their downstream effectors (red bars in **a**) are significant, and the interaction directions are consistent with the literature. **c.** Most of the top significant protein-protein interactions (blue bars in **a**) can be found as direct or indirect interactions in Pathway Commons (PC). The distributions of interaction strength across 1,000 models for each interaction in the two tables with corresponding colors are centered away from zero, in contrast to the background distributions of aggregated interactions across models (gray, all interactions with drug activity nodes in **a**, all protein-protein interactions in **c**). All other interactions with t-scores and PC information are included in Supplementary Table S2. **d.** A network visualization of the top interactions (top two interactions acting from each node and onto each node) highlights the level of agreement between model-inferred interactions and those in PC.

To further investigate to what extent the inferred networks represent known pathway interactions, we compared the interactions in the CellBox models to corresponding molecular interactions extracted from the Pathway Commons (PC) resource (Cerami *et al*., 2011), which is an aggregation of ∼20 curated publicly available pathway interaction databases. In this comparison between interactions inferred by CellBox models and prior knowledge, model interactions can, in principle, be found in PC either as one-step (A-B), two-step (A-X-B), or logical (>2 steps) interactions, where the logical interactions can be usefully predictive or perhaps erroneous. Direct interactions (A-B) in PC represent one component affecting the expression, phosphorylation, or state-of-change of the other component (B) (Figure 5c, 5d, Supplementary Table S2). For example, models and prior knowledge agree for the phosphorylation of the AKT protein kinase (AKTpS473) that negatively affects the insulin receptor substrate 1 (IRS1) (Chandarlapaty *et al*., 2011) and, for the activation of the mitogen-activated protein kinase (MAPKpT202) that affects p27 protein levels (Donovan, Milic and Slingerland, 2001; Osaki and Gama, 2013). Some model interactions can be found as indirect two-step interactions (A-X-B) in PC, meaning at least one path with one intermediate component(x) can be found between the two components (A and B). For example, retinoblastoma protein (Rb1) affects cyclinD through p21 (Carreira *et al*., 2005; Lei, Liu and Ness, 2005); and mitogen-activated protein kinase kinase (MEK1) indirectly interacts in two steps with the transcription factor c-Myc by a phosphorylation mechanism via ERK1/2 (Gupta and Davis, 1994; Butch and Guan, 1996; Sears, 2000; Aoki *et al*., 2011). Additional evidence of functional links between proteins identified in the models includes protein-protein edges in the STRING database (von Mering *et al*., 2005), such as links that represent expression (mRNA) correlation, and PubMed-derived links from text-mining (Maglott *et al*., 2011).

In order to confirm that such agreement between the inferred networks and prior knowledge is not an artifact, we examined solution stability as well as compared the inferred networks to random networks. We conducted stability selection tests (Meinshausen and Bühlmann, 2010) on the interaction parameters, and the results indicate the solutions are reproducible and robust (Figure S9a). We compared the inferred networks to networks with randomly drawn interactions (see Methods). In the inferred networks, we observed a statistically significant higher number of top interactions consistent with the knowledge in Pathway Commons (PC), indicating that such agreement is not an artifact of the densely connected signaling networks in the pathway database (Figure S9b). The remaining model interactions are between components that are more than two pathway steps away or cannot be connected by any path in PC. These interactions can be interpreted as either logical interactions important for predictive purposes, or potential new physical interactions that are yet to be discovered in molecular experiments. Also, the Pathway Commons (PC) database aggregates interaction information from multiple biological systems, and, therefore, complete consistency is not expected for any particular system, e.g., the melanoma cell line used here. Since the CellBox network models are constructed in a completely data-driven way without any a priori intention to recapitulate known molecular interactions, the partial agreement between the model-inferred molecular interactions and the experimentally known ones from the literature is evidence of the validity of the modeling approach.

### Predictions of unseen perturbations give candidates for drug combinations

Our results so far indicate that the CellBox models can be efficiently trained on a relatively small set of experimental data to parametrize the differential equations that model the behavior of the entire system of nodes and interactions at a reasonable level of predictive accuracy. This model can then predict cell responses to a full range of single and combinatorial unseen perturbations, which would be laborious and costly to test exhaustively by experiments. In order to nominate effective drug combinations for a much reduced number of focused experiments, we used simulations of the 1,000 full models to quantitatively predict the dynamic cell responses to ∼160,000 in silico perturbations, including different dosages of single perturbations on each protein node as well as all pairwise combinations (see Methods). For each perturbation condition, we averaged the predictions across all models and ranked the perturbations by predicted phenotypic changes (Figure 6a).

**Figure 6.**
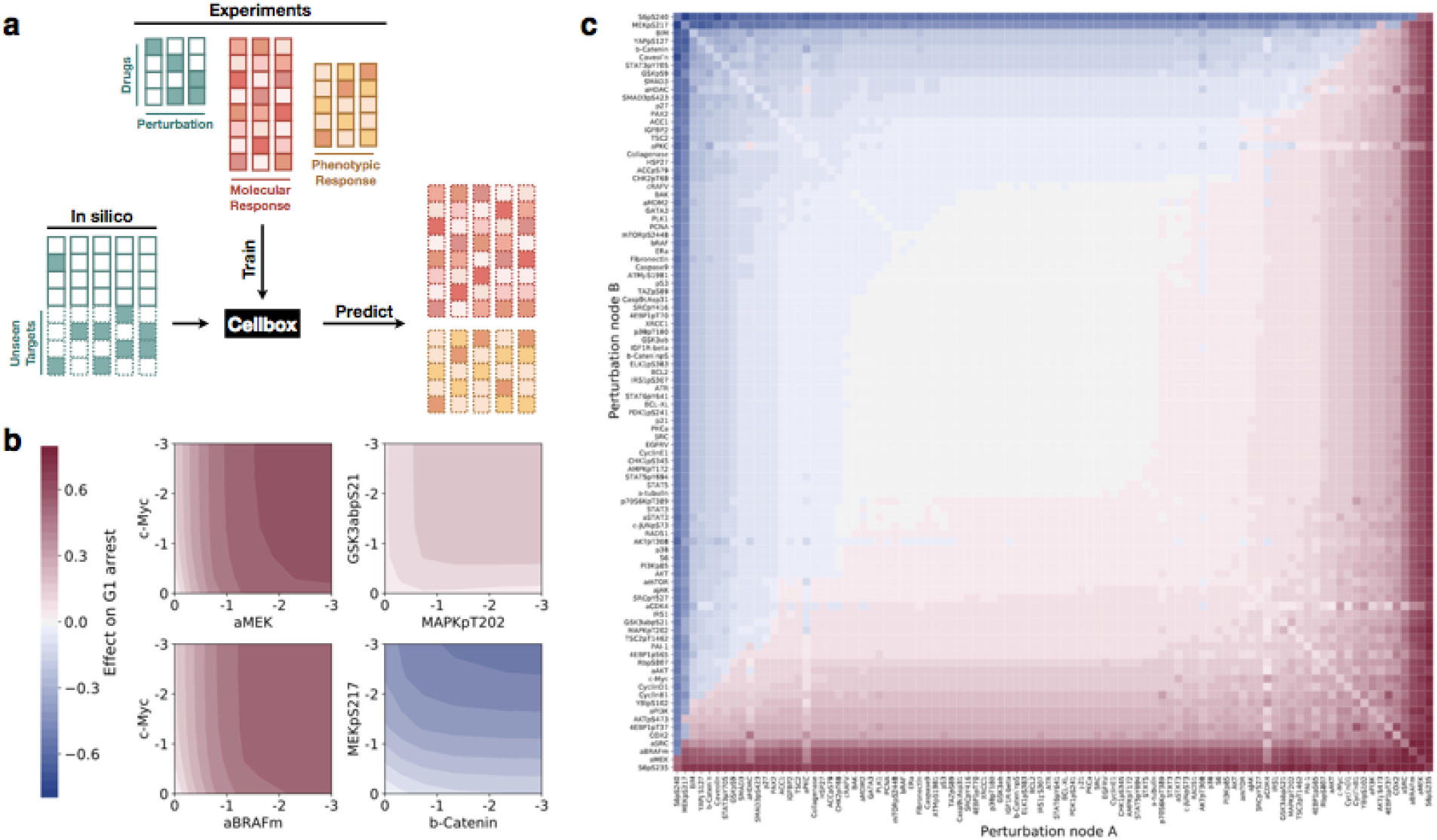
CellBox provides testable predictions of cell phenotype under synthetic perturbations. **a**. For each (phospho)protein node in the network, CellBox was used to simulate all single and paired inhibitions and to predict the phenotypic changes. The phenotypic effects are the average prediction of 1,000 independent models trained on the full datasets. **b**. The anti-proliferation effects of two perturbation pairs whose effects on cell cycle arrest have been experimentally tested were closely examined (left two panels, c-Myc+MEKi, and c-Myc+RAFi), as well as two other in silico conditions (right two panels, GSK3p+MAPKp and MEKp+b-Catenin), by simulating with combinatorial perturbation strengths. **c**. The effects on cell cycle arrest of pairwise combinatorial perturbation of all (phospho)proteins in the network were simulated and used to nominate effective pharmaceutical candidates. These in silico inhibitory perturbations can result in anti-proliferation effects (red, bottom right) or pro-proliferation effects (blue, top left).

Previous models on the same dataset, using the same differential equations but parametrized using belief propagation, had predicted that two drug pairs, MEKi+c-Myc and RAFi+c-Myc, would increase G1 cell cycle arrest and this prediction was confirmed by experiments (Korkut *et al*., 2015). We found that the CellBox model predicts similar effects for these two drug pairs (Figure 6b). In order to identify additional therapeutic candidates, we examined the effects of all possible single and pairwise perturbations on cell cycle arrest (Figure 6b, 6c). The top-ranked candidates included dominant anti-proliferative inhibition (uniform colors in rows or columns) of proteins in the Wnt, MAPK, and ERK/MEK pathways, known to be cancer-related. Besides strong single candidates, synergistic drug pairs are of potential therapeutic interest (Figure 6b, 6c departure from uniform colors). Inhibitory perturbations predicted to have pro-proliferation effects, which are undesirable as such, can also lead to effective anti-proliferative candidates via indirectly activating perturbations (Figure 6c, top left corner). For example, protein nodes can, in principle, be activated by reducing upstream inhibition or degradation. As the CellBox model is completely data-driven, the *de novo* predictions represent system-specific predictions independent of prior knowledge.

## Discussion

Quantitative models that are predictive of dynamic cellular responses can be used to design combination therapies in cancer. To provide predictions with sufficient accuracy and potential mechanistic insight, we integrated machine learning methods with dynamic modeling: we applied an optimization algorithm used in deep learning to a biologically interpretable differential equation (ODE) system. Our model CellBox can be trained efficiently and independently of prior knowledge to predict molecular and phenotypic responses to unseen perturbations with high accuracy. Although trained on a relatively small set of experiments, the model is capable of simulating cell responses to numerous arbitrary combinatorial perturbations and dosages applied to nodes repeatedly measured under different perturbation conditions. Ranking of cellular responses to the in silico combinatorial perturbations by the desired phenotypic outcome, such as decreased proliferation, potentially leads to specific therapeutic hypotheses.

Interpretability of models that are to be used for practical decisions, such as the design of combination therapy, helps increase confidence and facilitates the design of focused validation experiments and is therefore as important as accuracy (Lipton, 2018). CellBox features interpretability in two aspects: transparency and traceability. *Transparency*: by using a well-defined mathematical model, CellBox is designed to be explicitly interpretable. In the current ODE model, each parameter represents a directed and quantitative interaction between cellular components or phenotypic quantities. *Traceability*: given a perturbation to the cellular system, the ODE simulation indicates how the effects of the perturbation propagate throughout the directed network in a time-dependent manner. The models can, therefore, provide mechanistic hypotheses of how the perturbations cause the observable cellular responses. Our model aims to meet the emerging demands of explainable artificial intelligence (XAI), which is essential for a socially acceptable application of machine learning models in biomedicine (Thelisson, 2017; Gunning and Aha, 2019).

In principle, CellBox is generalizable to other types of systems and larger systems. Other types of models will presumably benefit from automatic differentiation (AD) combined with stochastic gradient descent that performs optimization directly for any given mathematical ansatz and, therefore, can avoid oversimplified approximations (Baydin *et al*., 2018). The flexible AD framework allows the models to be easily adapted to various forms of cellular kinetics and dynamics. The ability to model larger systems depends both on the availability of larger datasets and scalable modeling methods. Larger datasets can be obtained by measuring diverse types of molecular data, for example, transcriptomic, epigenomic, and metabolomic changes (Brown *et al*., 2014; Zaal and Berkers, 2018). A major opportunity for larger datasets may arise from recent cell barcoding techniques that significantly increase perturbation throughput relative to arrayed experiments (Adamson *et al*., 2016; Dixit *et al*., 2016) by measuring levels of transcripts by sequencing, levels of proteins detected by antibodies labeled by oligonucleotides or isotopes at the single cell level, or levels of both by multiplexing (Frei *et al*., 2016; Wroblewska *et al*., 2018; Mimitou *et al*., 2019; Schraivogel *et al*., 2020). A key advantage of single cell perturbation approaches for data-driven inference of dynamic networks is scale, as barcoded perturbation experiments can be pooled and individual cells identified based on sequence tags.

As CellBox is implemented in the Google TensorFlow framework, it can make use of various advanced machine learning techniques, such as dropout, mini-batching, and GPU boosting (Bengio, 2012; Liang and Liu, 2015), to improve training efficiency, which partially addresses the issue of scalability (Stapor *et al*., 2019). As most state-of-art single-cell sequencing technologies still suffer from limited sequencing depth and inevitable noise, a new set of computational challenges arises when applying CellBox models to Perturb-seq like datasets. Future efforts are needed for resolving such issues of sparsity and stochasticity, e.g., using robust techniques of dimensionality reduction (Lopez *et al*., 2018; Eraslan *et al*., 2019; Lotfollahi, Wolf and Theis, 2019).

A particular advantage of our modeling approach is its complete independence from prior knowledge. This is in contrast, e.g., to other frameworks for interpretable models (Fröhlich *et al*., 2018) that incorporate prior knowledge by pre-defining the connections in the network model based on large-scale curation of molecular and biochemical pathways. Such models have been further developed and incorporated with proteome/phosphoproteome data (Schmiester *et al*., 2020). While our current implementation of the model is completely data-driven, prior knowledge of cellular interactions could also be included by adding a penalty to the optimization function, which quantifies the disagreement between inferred and prior values for each parameter. However, considering the incompleteness of pathway knowledge bases and relatively high prediction accuracy of CellBox, we speculate that applying prior knowledge might give limited improvement in model performance. Besides, it would also be of interest to explore whether one can combine the efficiency advantage in Fröhlich et al. (Fröhlich *et al*., 2018) via adjoint analysis (Fröhlich *et al*., 2017) with the advantages of the current implementation of CellBox.

For broad translational applicability, a tantalizing but challenging prospect is to incorporate individual tumor background via genetic perturbations and to propose optimal, personalized combinations of targeted therapeutics. We envision the systems biology approach described here to be broadly applicable to other areas of biology, such as developmental biology or synthetic biology, provided that suitable perturbation-response data becomes available. Key future challenges are, therefore, the design of experiments for each biological context of interest and the further development of transferable and scalable machine learning methods.

## Methods

### Perturbation dataset overview

The CellBox models were trained using a perturbation-response dataset of the SK-Mel-133 melanoma cell line (Korkut *et al*., 2015) (Figure S1). The cells were treated with 12 different single drugs, each at two different concentrations and 66 pairwise combinations of these drugs at IC40 concentrations. 24 hours after drug treatment, Reverse Phase Protein Arrays (RPPA) wasused to measure the level of 45 proteins and 37 phosphoproteins of interest. Cell cycle progression, including G1 arrest, G2 arrest, G2/M transition, and S arrest, was measured by flow cytometry. Cell viability was measured 72 hours after drug treatment by the resazurin assay. Our experimental data was normalized using median normalization, which is a standard approach of processing RPPA data (Tibes *et al*., 2006). We used the log_2_ ratio of measurements in perturbed conditions over unperturbed conditions as the system variables *x_i_* in the models. The dataset was initialized with 12 drug activity nodes representing the inhibition strengths of different drugs to their targets (Molinelli *et al*., 2013). The resulting dataset has 89 perturbation conditions and 99 observed nodes (82 protein and phosphoproteins, 5 phenotypes, and 12 drug activity). A more detailed description of the experimental dataset is available in Korkut et al. (Korkut *et al*., 2015)

### Model configuration

The models were constructed using Python 3.6 and Google Tensorflow (Martín Abadi *et al*., 2015) (version = 1.15.0). The molecular and phenotypic changes are linked in a unified biological network model using a system of ordinary differential equations

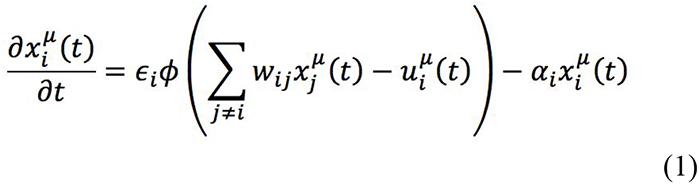

Where 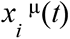 represents the log_2_-normalized relative change of each (phospho)protein or phenotype levels relative to control levels under condition μ. *u_i_*^μ^(*t*) quantifies the strength of the perturbation on target (*i*). Here the drug effect is assumed to be constant and, therefore, *u*(*t*) = *u* for *t* > *t*_0_ is the perturbation strength determined using the endpoint level change of the primary target of the particular drug (Figure S1b). α*_i_* characterizes the effect of decay, meaning the tendency of protein *i* to return to the original level before perturbation. The interaction parameters *^w^ij* indicate interactions between network node *j* on network node *i*, assumed to be a constant property of the pair of molecules in this given cellular setting. We constrain the interaction parameters *^w^ij* by disallowing three classes of interactions:

1. ingoing connections for drug nodes (drugs cannot be acted upon by any other node)
2. outgoing connections for phenotypic nodes (phenotypes cannot act on any other nodes)
3. self-interaction (nodes cannot act on themselves)

We use a sigmoid function ϕ(*x*) = *tanh*(*x*), to introduce a saturation effect of the interaction term so that it is bounded by the constant value of ε*_i_*. Imposing bounds on the absolute value of the contribution of the interaction term to the derivatives is in part motivated by the fact that we are only modeling a small fraction of cellular components and by the experimental observation of system stability in these experiments. We also tested different alternatives of envelope forms, including a clipped linear function (hard tanh) ϕ(*x*) = *max*(− 1, *min*(1, *x*)), symmetric polynomial function 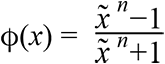, *x*′ = *max*(0, *x*) (adapted from Hill’s equation), and no envelope function ϕ(*x*) = *x* (linear) (Figure S5).

The biological network interactions were constructed *de novo* without any prior knowledge input, meaning the network was fully connected with interaction parameters.

The interaction parameters *^w^ij* were randomly initialized following a normal distributioñ *N* (0.01, 1). The other two coefficients α*_i_* and ε*_i_* were initialized as 1.0.

Taken together, with this formulation of system dynamics, the effect of each new input perturbation is quantified by the response of downstream protein and phenotypic nodes, and the effect of the perturbation is simulated by propagating the input across the inferred interaction network according to the differential equations. After learning from perturbation-response data, the model can therefore make predictions for responses of perturbations on any experimentally probed node, including nodes not perturbed in the experimental dataset from which the models are derived. The ODE system was numerically solved using Heun’s method (Süli and Mayers, 2003) (eqn. 2, time steps *N_t_* = 400, supplementary Figure S2), which is an improved variant of Euler’s method. Model performance was evaluated by disagreement between the experimental cell responses and the numerical steady state levels.

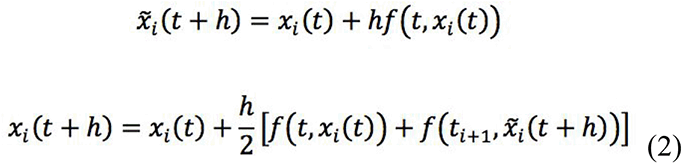

Where *h* is the step size, *f*(*t*, *x*(*t*)) = *x*’(*t*), *y*(*t*_0_) = log(1) = 0.

The loss function *L* (*w*) is defined as a weighted sum of prediction error and complexity penalty in order to avoid overfitting. Here a mean squared error (MSE) and an L1-loss regularization term are used, as defined in eqn. 3. We have tested different regularizations, including L1, L2, and both combined (elastic net) with various strengths (regularization strengths for L1, L2 term λ_1_, λ_2_ = 0, 0.01, 0.0001, pairwise combination), and found no significant difference in model performance (Figure S2b). The interaction parameters were optimized end-to-end using the Adam optimizer (Kingma and Ba, 2014), with the objective of minimizing the loss function.

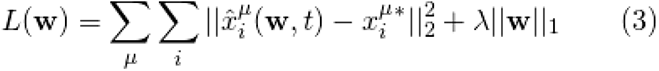

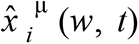 is calculated as the converged value of the numerical simulation of the ODE with the interaction parameters *w* and defined simulation timestep 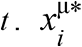 indicates experimental measurement which is used as a gold standard in training. The dataset was divided into training, validation, and test sets, in order to optimize parameters, provide an indication for stopping training, and test model performance, respectively. Optimization was conducted with an initial learning rate for the Adam optimizer (lr=0.1) and regularization strength (λ =0.01). It has been shown that a gradually decreasing learning rate is helpful for model convergence (Bengio, 2012). The model training was stopped when the loss function of the validation set does not further decrease for a continuous of 20 iterations (stopping patience).

The model was trained with mini-batching: a random 80% portion of the training set was used to optimize parameters for each iteration, and a fixed batch size of 4 was used in each epoch. Models that failed to converge (MSE for training set > 0.05) were excluded as unsuitable. For a larger dataset, we recommend using a fixed batch size of 8 or 16, as documented in the latest version of the software on GitHub.

### Model training with random data partitions

For initial model training and analysis of model performance, the cell line perturbation-response dataset was randomly partitioned into training, validation, and test set in the proportion of 56% (n=50 conditions), 14% (n=12 conditions), and 30% (n=27 conditions). 1,500 models were generated on 1,500 independently random-partitioned datasets.

The models were examined and categorized into non-oscillating and oscillating solutions based on time derivatives at the final time step of the ODE simulation. The non-oscillating solutions are defined as those with the average absolute value of time derivatives of all nodes and conditions in the training set smaller than δ, i.e. 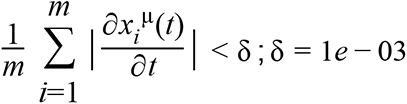. In each category, twenty models were randomly selected, and each re-trained with the original data partitioning but forty different random seeds, covering all the random processes in training, including parameter initialization and mini-batching sampling (Figure S4). Oscillating solutions comprise about 30 percent of all models. In the following analysis, models that converged to oscillating solutions were excluded.

Under the training scheme using random data partition, we examined the sensitivity to data scaling and to the choice of ODE solver, by applying the following modifications to the data or the model and evaluating the changes of model performance. i) In addition to the log_2_ scale of the ratio of measurements in perturbed conditions over unperturbed conditions, we applied other scalings to the raw data to generate model input, including non-log (linear) scale (Figure S3c). ii) In addition to Heun’s ODE solver, we tested other numerical ODE solvers, including Euler methods, Midpoint methods, and Runge-Kutta methods (Figure S4e-g).

### Single-to-combo and leave-one-drug-out cross-validations

In the single-to-combo task, all single-drug conditions were allocated to the training set (n=23 conditions), and the combination perturbation conditions were randomly distributed among the validation and test set (n=53 conditions) in a 20/80 ratio. To evaluate model performance by cross-validation for each drug, the data was partitioned into training (n=78 conditions) and test (n=11 conditions) sets, where each test set contains all the drug combination conditions with the particular drug. 20% of training conditions (n=15) are used as a validation set. The predictions on the test set were averaged over 100 models. In the more stringent leave-one-drug-out task where all perturbation conditions involving one drug were withheld as the test set, the evaluation of model performance was conducted by applying perturbation of the same strength directly on the downstream effector node of each withheld drug activity node (Figure S1b).

The Belief Propagation (BP) models for both single-to-combo and cross-validation prediction were performed as in our earlier publication (https://github.com/korkutlab/pertbio). The predictions on the test set were averaged over 100 models. The deep neural network model (NN) network had 2 densely connected hidden layers (hidden layer H1: 20 neurons, H2: 100 neurons), connecting the parameterized perturbation tensor with the cell response tensor. We used the *tanh* activation function for the hidden layers and the leaky rectified linear unit function (ReLU) for the output layer. The NN models were constructed in the Tensorflow framework in Python and optimized using the same optimization methods (Adam optimizer). The co-expression static model (Co-exp) was constructed in the Python environment using the sklearn (version = 0.21.3, https://scikit-learn.org/stable/) *MultiTaskElasticNetCV* module, which is a linear regression model with L1 and L2 regularizations whose strengths are automatically determined using built-in cross-validation. The model was trained to use the changes in levels of each pair of protein nodes (input) to predict levels for the rest of the proteins and phenotypes (output). To compare the four different predictive models, CellBox, BP, NN, and Co-exp, the paired t-test on two related samples was used to analyze the difference between model performance (test set correlation) and to assign p-values.

### Sensitivity analysis with noise and reduced training set size

We conducted a sensitivity analysis of our model to evaluate the robustness of its prediction in response to noise. To model experimental noise, we first applied varying levels of multiplicative Gaussian noise to the input molecular and phenotypic data (eqn. 4). The assumption is that the uncertainty and noise in experimental measurements arise around the true values with a scaling factor depending on the experimental approach.

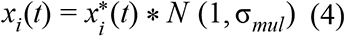

The scaling factor for each node and each condition was independently drawn from a Gaussian distribution *N* (1, σ*_mul_*), with a mean of 1 and a standard deviation of 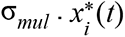 represents the experimental measurements, and represents the values with noise added. For each noise level, we evaluated 15 different training/validation/test partitioning and each with 5 independent random noise patterns. Model training was performed on noisy training and validation sets, while the model performance evaluation was performed on the original, noise-free test data. For each noise level, the percentage of successful models, defined as those that converged in terms of both MSE and oscillation filters, was recorded.

To further evaluate model performance from a technical perspective, we applied additive Gaussian noise to the input data (eqn. 5) and evaluated model performance as above.

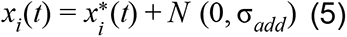

We examined model sensitivity to training and validation set size. We reduced the combined size of the training and validation set, from 90% to 10% in steps of 10%, while keeping their relative size constant, 4:1. The remaining data were allocated to the test set. For each training set size, the percentage of successful models, defined as those that converged in terms of both MSE and oscillation filters, was reported.

### Biological interpretation of the network model

The entire dataset was used to generate 1,000 full successful models, each with an independent data partitioning of training (n=71 conditions) and validation (n=18 conditions). For each interaction (*w_ij_*) between two nodes, a t-score 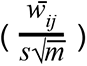 (which is effectively an one-sample t-test with the null hypothesis that the population mean is zero) was calculated as an indication of the confidence level of obtaining a value different from zero, where 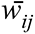 is the average interaction strength across models, *s* is the standard deviation, *m* is the number of models (m = 1,000) (Supplementary Table S2).

In order to compare the model inferred interactions to those present in prior-knowledge pathway databases, all the proteins and phosphoproteins nodes were identified by their corresponding gene names (Supplementary Table S1). The interactions were compared against the Pathway Commons (PC) database (current version at https://www.pathwaycommons.org/archives/PC2/v11) using the paxtoolsr (Luna *et al*., 2016) software. The database was filtered down to direct interactions, which include “controls expression of”, “controls phosphorylation of”, “controls state change of”, “controls production of”, “controls transport of”, and “controls transport of chemical”. All the direct interactions were converted to a directed PC graph using igraph (version = 1.2.4.1, https://igraph.org/r/). Distance was calculated as the length of the shortest path (s) between two components in the PC graph (Supplementary Table S2, column: distance). The number of the shortest paths (npath) between the two components was also reported. Additional detailed information on protein links in Homo sapiens was obtained from the STRING database, including combined score (string) and individual-channel scores (co-expression, experimental, database, text-mining) (https://stringdb-static.org/download/protein.links.detailed.v11.0/9606.protein.links.detailed.v11.0.txt.gz) (von Mering *et al*., 2005). The co-citation score was calculated as the number of papers mentioning both components in a customized paper collection aggregating all PubMed papers that refer to at least two and no more than five gene names (Maglott *et al*., 2011).

To quantitatively evaluate whether the inferred networks recapitulate more known interactions in the database than the random networks, the same number of network models (m = 1,000) were generated with interaction parameters randomly drawn from the pool of all (phospho)protein-(phospho)protein interaction parameters in the CellBox-inferred network models. For each of the inferred or random network models, the number of interactions in Pathway Commons (PC) out of the top one-hundred interactions (ranked by absolute interaction strengths) was calculated and the t-test was performed to compare the difference in means between the two groups. To provide an additional comparison with a well-known alternative method for network inference, the partial information decomposition and context (PIDC) algorithm was implemented in the Python environment based on the detailed mathematical description of the algorithm in Chan et al (Chan, Stumpf and Babtie, 2017). The same number of PIDC networks (m = 1,000, same as for CellBox) were generated based on 1,000 independent random partitions of the entire dataset (80% of the conditions were used to infer PIDC networks). The t-test was performed to compare the difference in means between the number of interactions inferred by CellBox network models and PIDC networks consistent with the PC database (Figure S9).

### Model predictions for a large number of unseen perturbations

We used the models trained with the full (non-partitioned) dataset to simulate responses of novel, experimentally unobserved, in silico perturbation conditions. These conditions included different doses of single perturbations (different levels of perturbation strength *u* ∈ [0, 3] on all individual (phospho)protein and drug activity nodes within the network and as well as all pairwise combinations 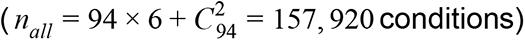. The cell responses were dynamically simulated with u as the input perturbation with the same number of steps as in training (*N_t_* = 400). For each perturbation condition, predictions for cell responses were averaged across 1,000 different models. Perturbations were nominated as therapeutic candidates by ranking the predicted magnitude of the phenotypic change in terms of cell cycle arrest.

### Code and data availability

CellBox code and data are available at https://github.com/dfci/CellBox as release v0.2. We provide a shareable, online, interactive environment using Binder with all necessary dependencies pre-installed. The Quick Start section provides instructions to quickly try an example script that runs a shortened training process.

## Acknowledgements

We thank Alexandra Franz, Frank Poelwijk, Laura Kleiman, Nicholas Gauthier, Haozhe Shan, William Yuan, Han Altae-Tran and members of the Sander and Marks labs for constructive discussions. Funding support from DFCI, NHGRI (U41HG006623) and NIGMS (P41GM103504).

## Supplementary materials

**Figure S1.**
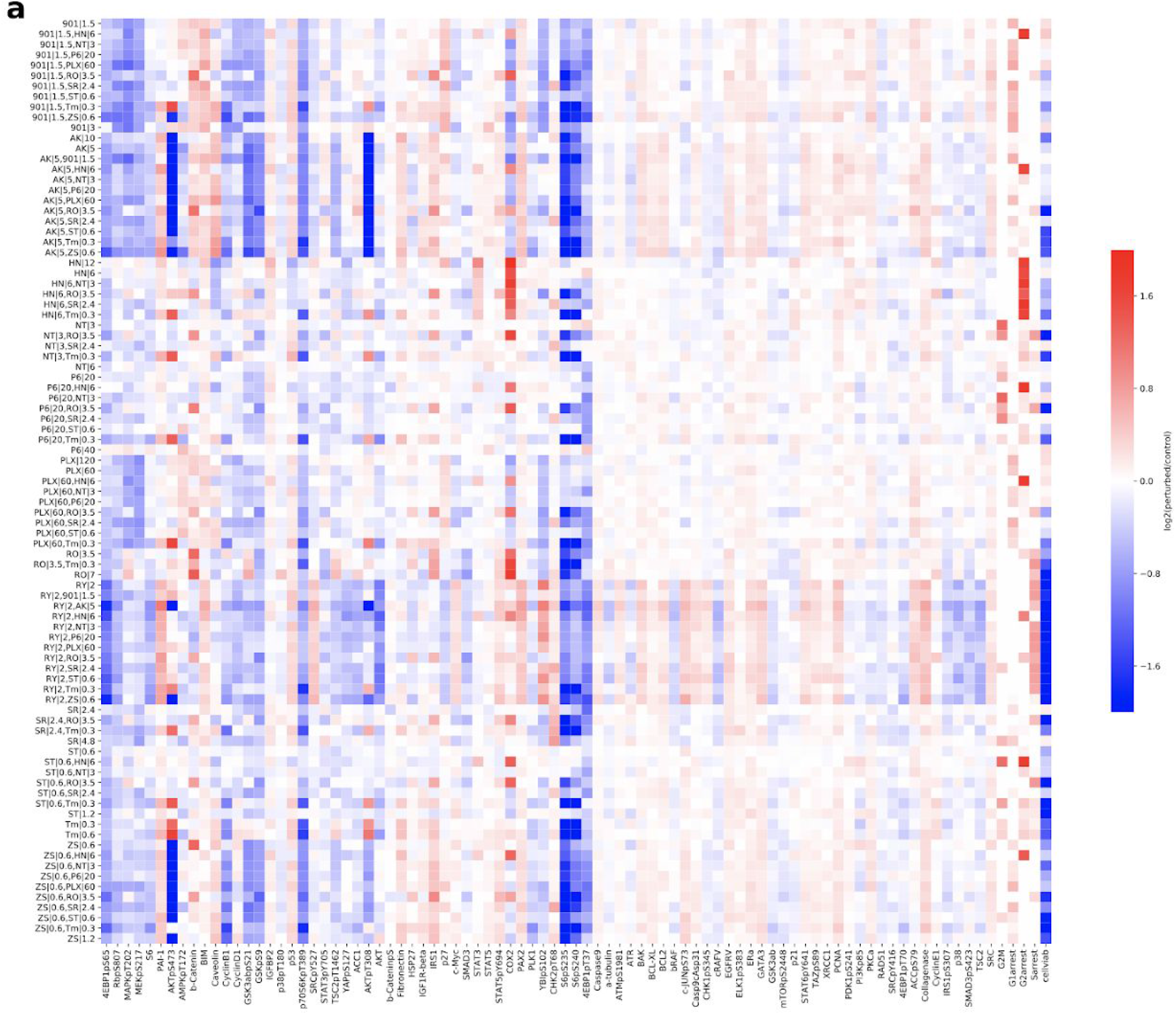
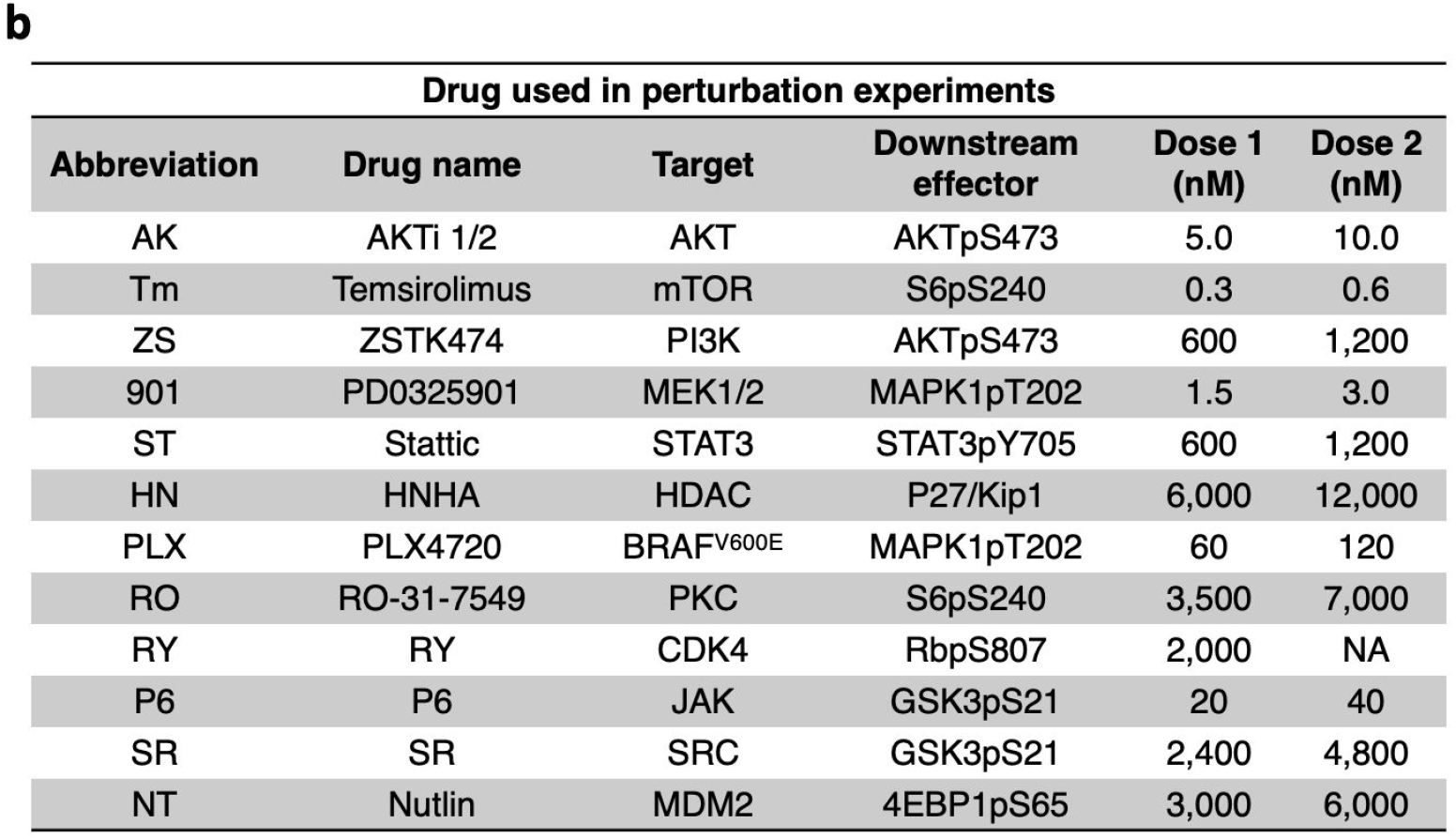
An overview of the perturbation dataset. **a.** The perturbation dataset (log_2_ transformation of the ratio of measurements in perturbed conditions over unperturbed condition) was visualized in a heatmap with rows of perturbation conditions (format: drug abbreviation with concentration, single or pair of drugs) and columns of (phospho)proteins or phenotypic measurements. **b.** 12 drugs were used in the perturbation experiments, and the table contains for each drug their abbreviations, targets, downstream effectors used in the inference of drug activity nodes, and the two doses used in single-drug perturbation conditions. In combinatorial perturbation conditions, the lower dose of each drug was combined.

**Figure S2.**
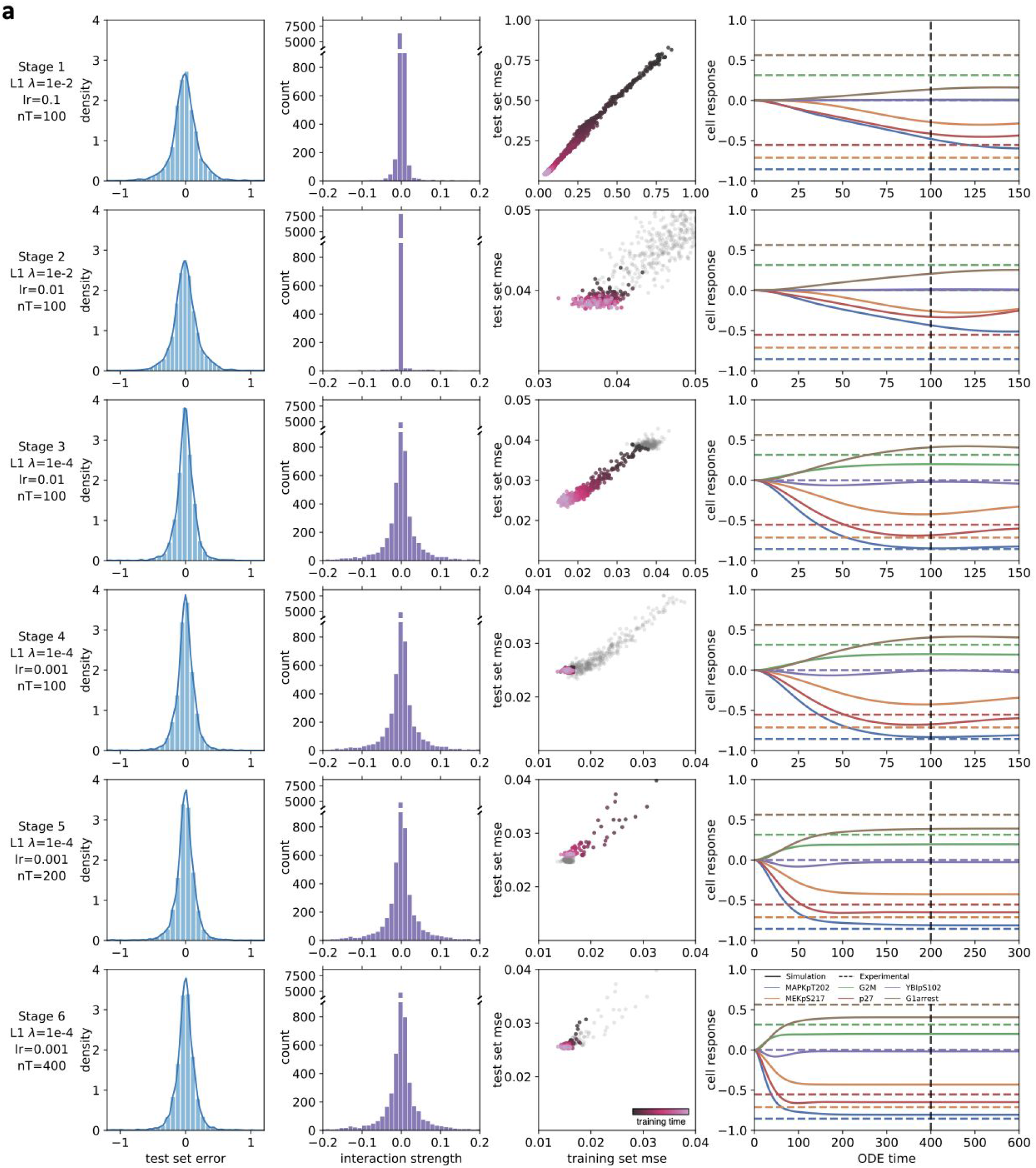
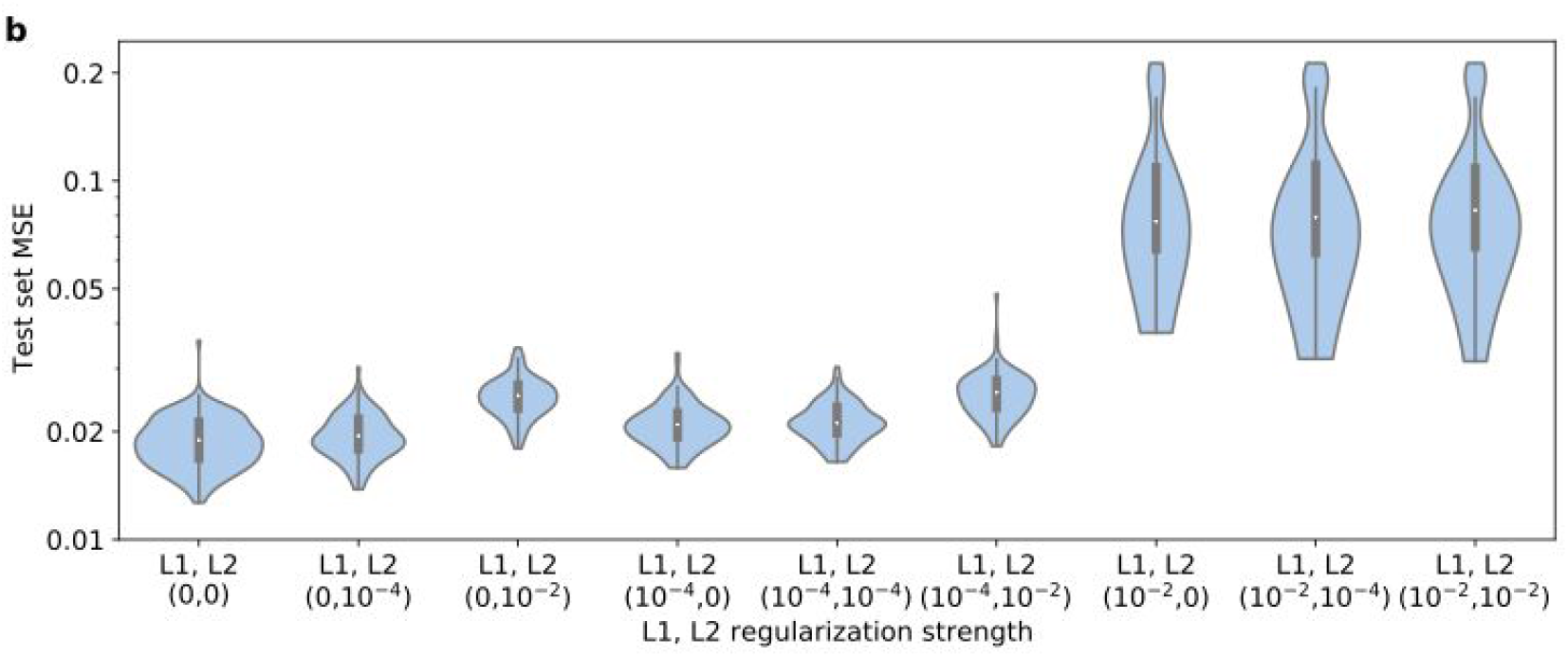
Multi-step fine-tuning of hyperparameters facilitates model training. During the training of each model, the training loss decreased along training time together with the test loss (Stage 1). The model parameters started to fluctuate around a local optimum after N = 2,000 training iterations. It has been shown that a decreasing learning rate is helpful for model convergence (Bengio, 2012). Decreasing the learning rate to 0.1x allowed the model to escape local minima and continue learning (Stage 2). The loss function stopped decreasing again after (up to) N = 4,000 iterations when the magnitude of the MSE loss was comparable to that of the regularization loss. To further improve training and decrease MSE loss mainly, we decreased the regularization strength by loosening the L1 constraints on the parameters (Stage 3). MSE decreased further while the numerical range of the parameters (interaction strengths) started to increase. Continuous decreasing of the learning rate did not further change the loss (Stage 4). ODE simulation of the model indicated that a steady state had not been fully reached. Therefore, the ODE simulation time was doubled twice (Stage 5 and Stage 6) while the learning rate and regularization were kept the same as Stage 4. The training and testing loss, together with the ODE trajectory, indicated that, at the final stage, the models had converged, optimization made no further improvements, and the ODE simulation has reached a steady state. We then stopped the training and examined the results closely on the test dataset. **a.** Models were trained in six individual stages with varying learning rate, regularization strength, and ODE simulation time. As the training proceeds, 1. the distribution of differences between predicted and experimental values in the test set narrowed around zero; 2. the distribution of interaction strengths widened as the L1 regularization was weakened; 3. both the training and test loss decreased; 4. ODE simulation reached a steady state as simulation time increased. **b.** Model performance was evaluated on different regularization approaches, including L1, L2, or combined (elastic net) with different regularization strengths (regularization strengths for L1, L2 term λ_1_, λ_2_ =0, 1e-2, 1e-4, pairwise combination). For a mild level of regularization (λ_1_ = 1e-4 and λ_2_ = 1e-4), we observed no significant difference between L1, L2, and elastic net. A stronger L1 regularization compromises the accuracy of model predictions.

**Figure S3.**
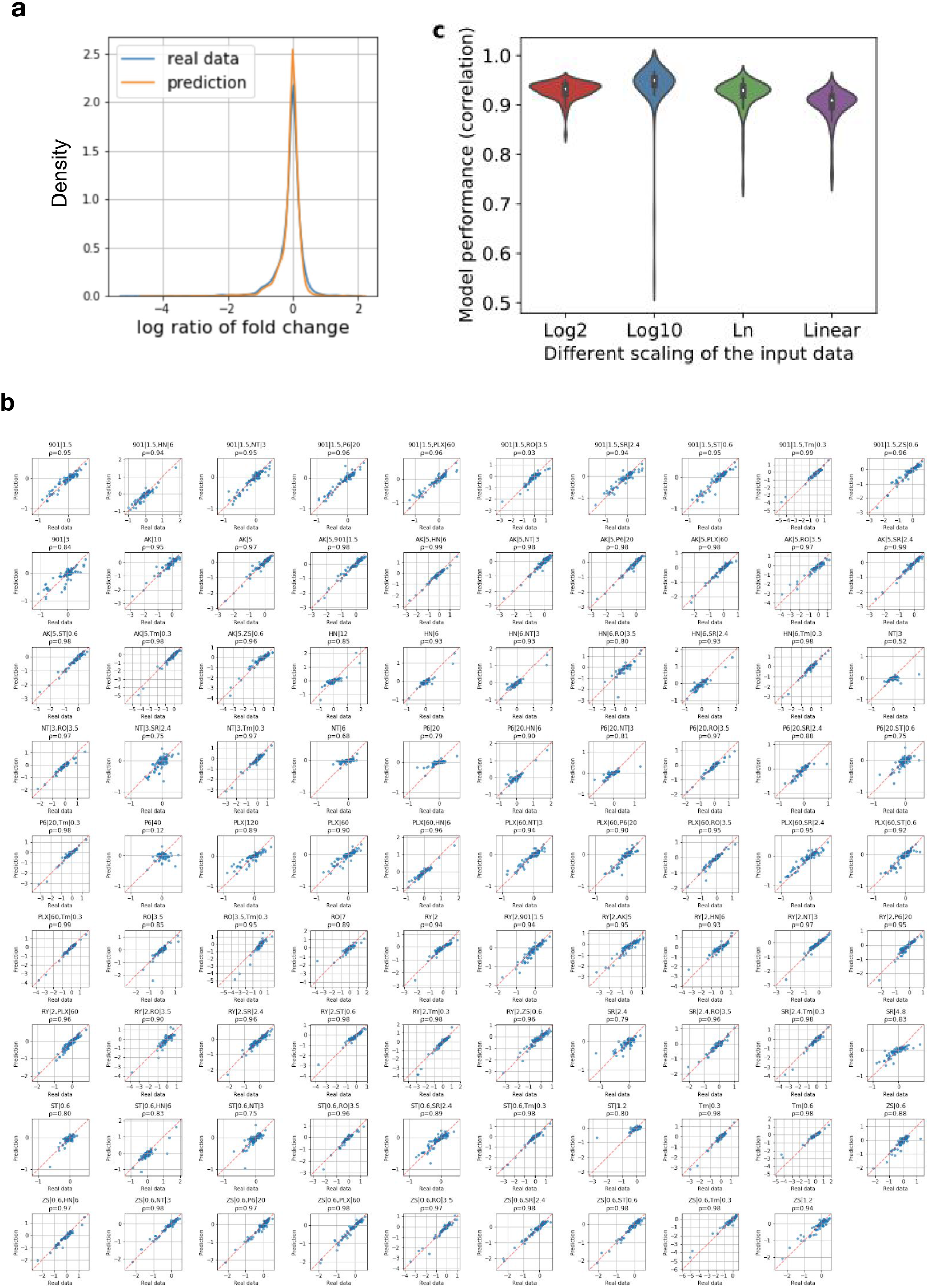
Correlations between predicted and experimental data were consistently high across different perturbation conditions. **a.** Model prediction and experimental data had a similar range and distribution without skewing and extreme predictions. **b**. In addition to overall performance, the model predictions for each perturbation condition were examined. The prediction of cell response under each condition reached similarly high correlation (median Pearson’s correlation 0.95) with experimental data. Meshes indicate the range of real data. Models generally performed better for conditions with larger data range. **c.** After median normalization of the RPPA data, the log_2_ ratio of measurements in perturbed conditions over unperturbed conditions was calculated and used as the input for the model. To evaluate whether the model performance depends on our choice of logarithmic scaling, we tested the model robustness against different scaling, including log_10_, natural log (ln), and non-log (linear) scaling. The model performance remains similar for different logarithmic bases (Pearson’s correlation coefficient for test set ρ>0.93). Without logarithmic transformation, the model training still converges and results in a slightly reduced predictivity (ρ=0.90). The performance of CellBox is robust against different scaling methods.

**Figure S4.**
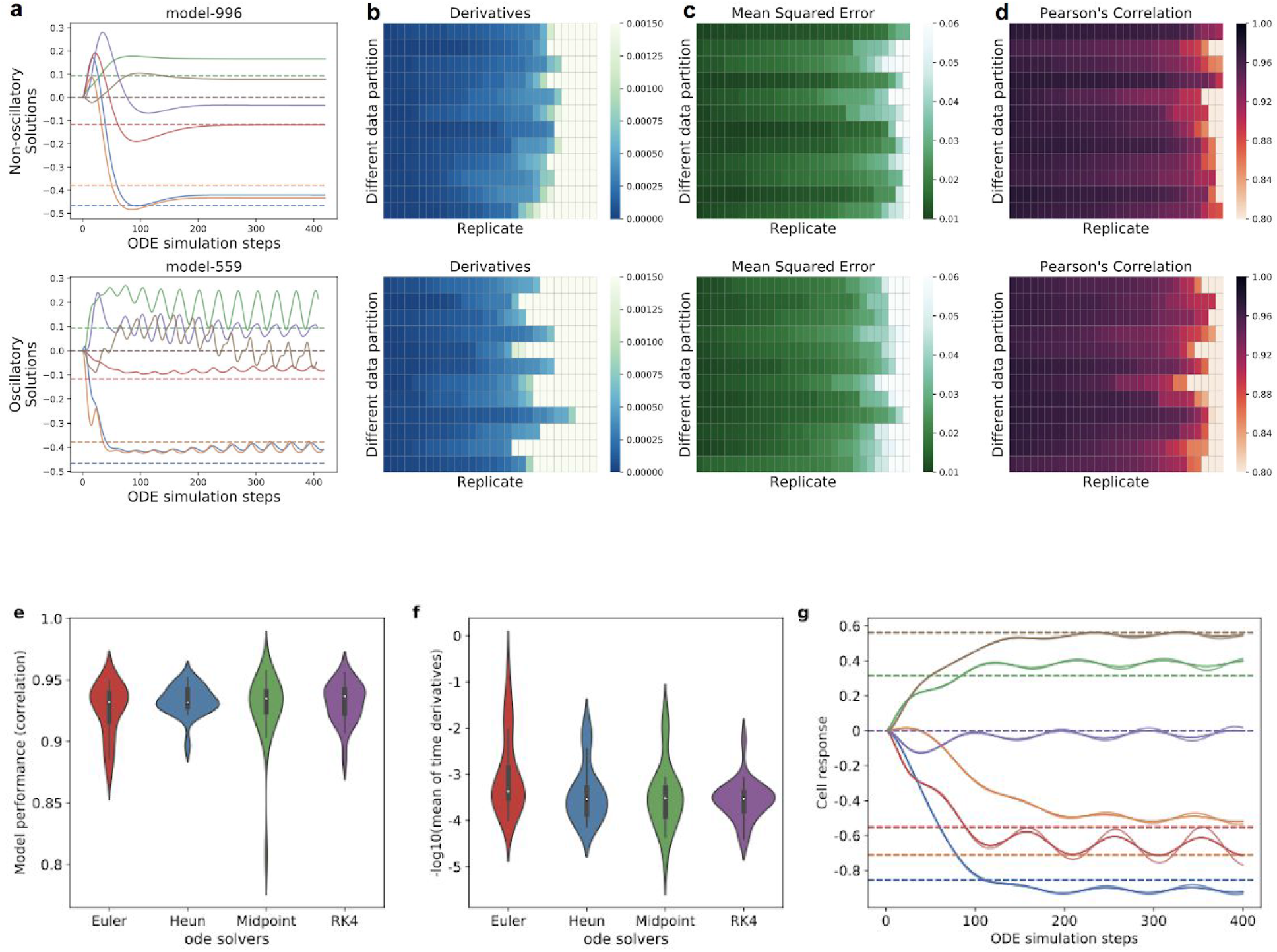
Oscillatory models from stochastic training are independent of data partitioning. **a.** Models were examined and categorized into non-oscillatory and oscillatory solutions based on the ODE simulation trajectories. **b-d.** For each data partitioning of training and test set (rows) in the two categories, different seeds for random processes in model training (columns). The models were examined for their performance in terms of average derivatives of each variable at the end of the ODE simulation (**b**), average mean squared error in the training set (**c**), and Pearson’s correlation between prediction and experimental data (**d**). Therefore, the solution oscillation and the model convergence are independent of the data partition. **e.** The model performance in terms of Pearson’s correlation between prediction and experimental data remains similar when different ODE solvers, including Euler, Heun, Midpoint and Runge-Kutta (RK4), were used in CellBox models. **f.** Oscillatory solutions (mean of time derivatives threshold δ = 1*e* − 03) exist when all different ODE solvers are used. **g.** For a given solution, all solvers are consistent in terms of whether the solution is oscillatory or not but might differ slightly in terms of the oscillation amplitude.

**Figure S5.**
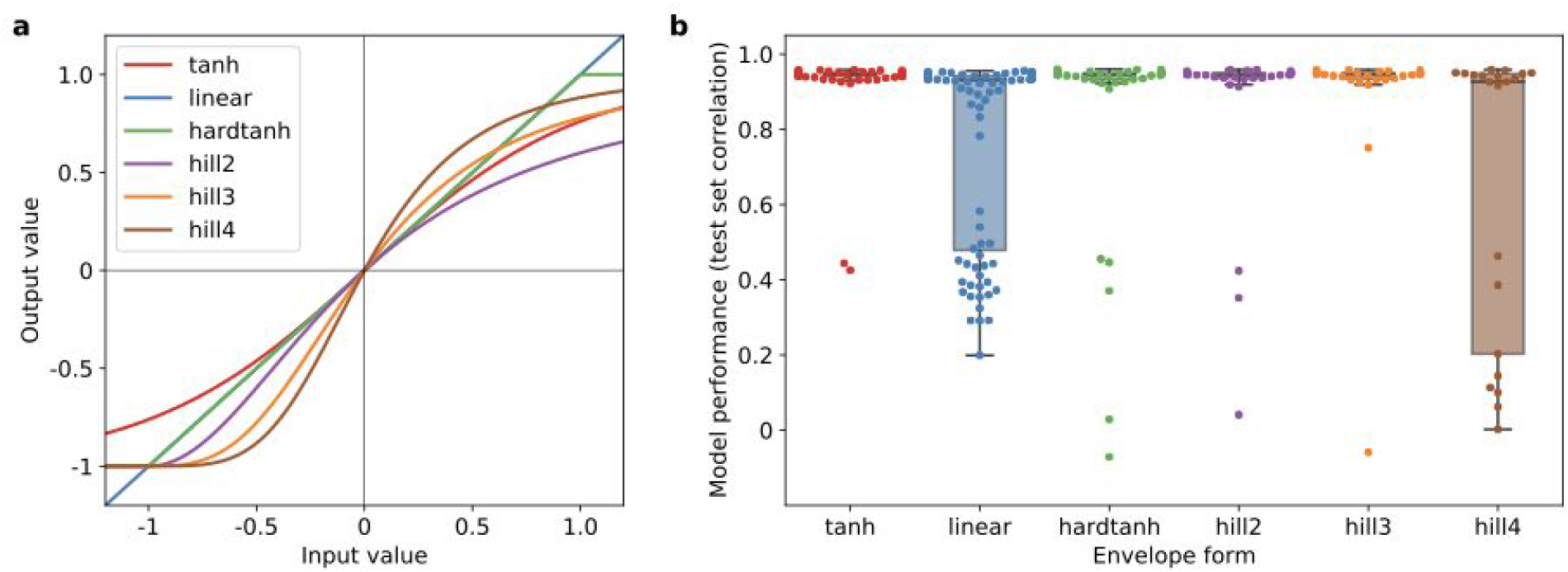
Model performance with different envelope forms. **a**. We tested different alternatives of envelope forms, including a hyperbolic *tanh* (used as current default in CellBox), a clipped linear function (hard *tanh*), symmetric sigmoid function (hill of various degrees), and no envelope function (linear). All the other envelope forms model the saturation effect except for the linear function. **b**. We observed comparable prediction accuracy between *tanh*, clipped linear, and low order of sigmoid functions, while the training with a linear envelope and high order of sigmoid functions was much less stable, and prediction performance was less accurate on average.

**Figure S6.**
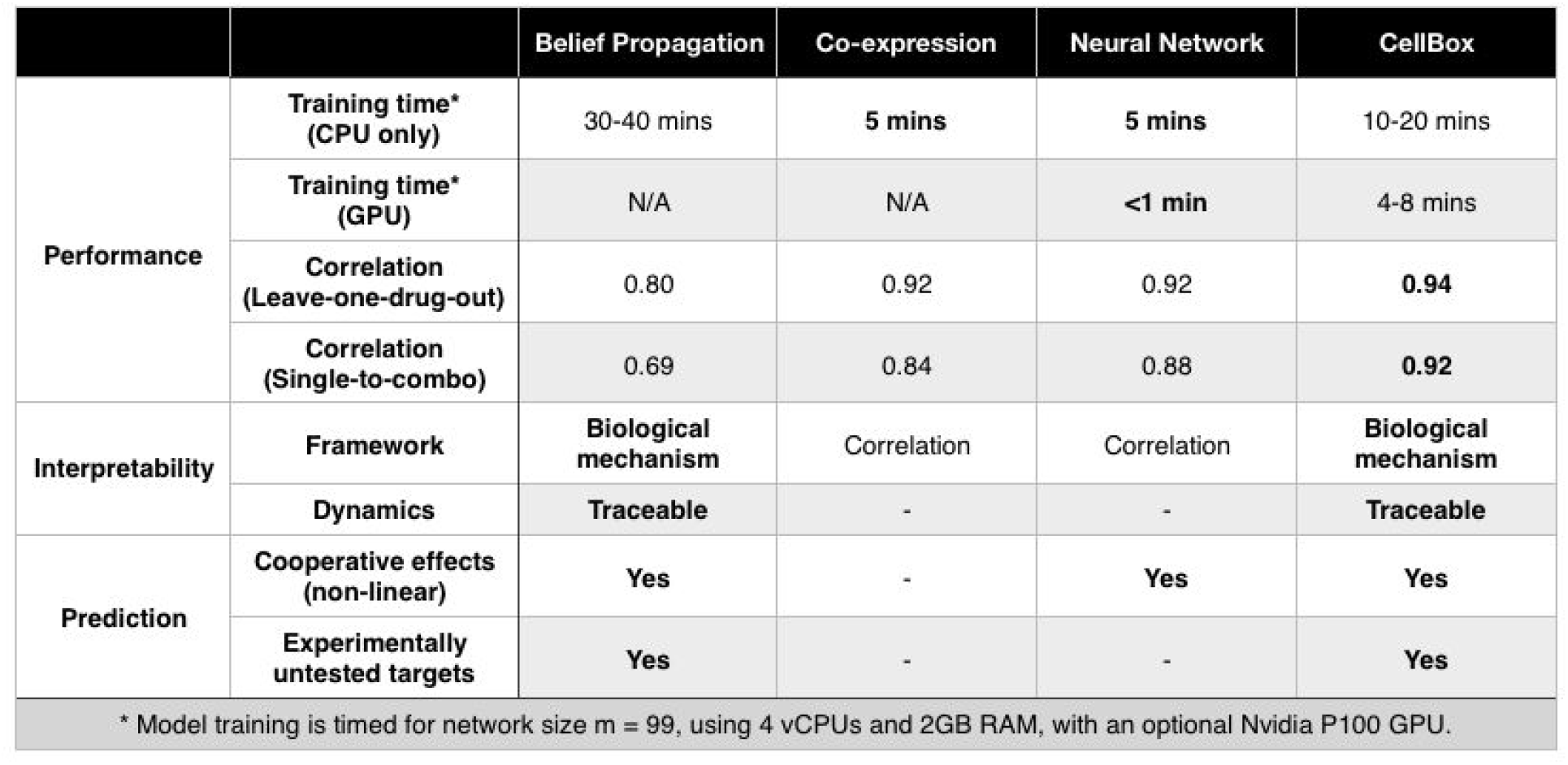
Comparison of various features between CellBox and other models. In addition to the comparison of model performance in several training schemes between different models (Figure 3), we provided a more detailed comparison between models in the perspectives of training time and model capabilities and labeled the best features out of all models (bold). Generally, we believe the faster the training is with a similar number of inferred parameters, and the more biological mechanisms the model can provide, the more generalizable and useful the model is.

**Figure S7.**
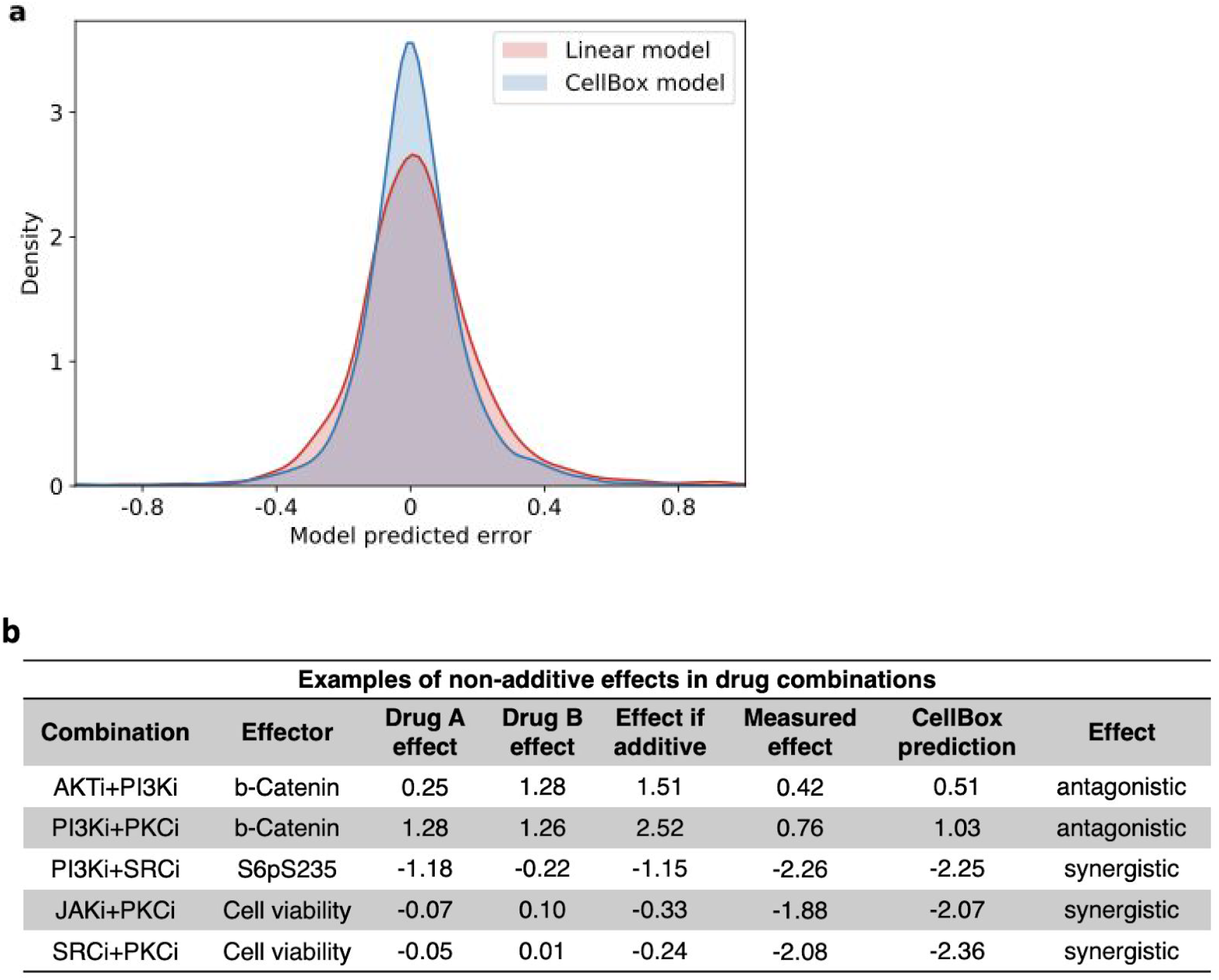
CellBox predicts the non-additive effects of drug combinations. One of the most important applications of CellBox is to raise candidates of synergistic drug combinations. In order to test model prediction for synergy, we examined the model predictions in the single-to-combo training scheme where only the single-drug perturbation conditions were used to predict cell response to drug combinations. We compared those predictions with additive predictions (linear model) and observed CellBox models could predict both synergistic and antagonistic effects. **a**. A systematic comparison of the prediction from CellBox and linear additive models. The CellBox models predict with higher accuracy (smaller prediction errors). The error is defined as the difference between model predictions and experimental responses for each data point. **b**. Examples of non-additive effects were chosen from the top ranking list of difference between prediction error from linear model and that from CellBox models, which are effectively the proteins or phenotypes in drug combinations that CellBox predicts accurately while the linear model does not.

**Figure S8.**
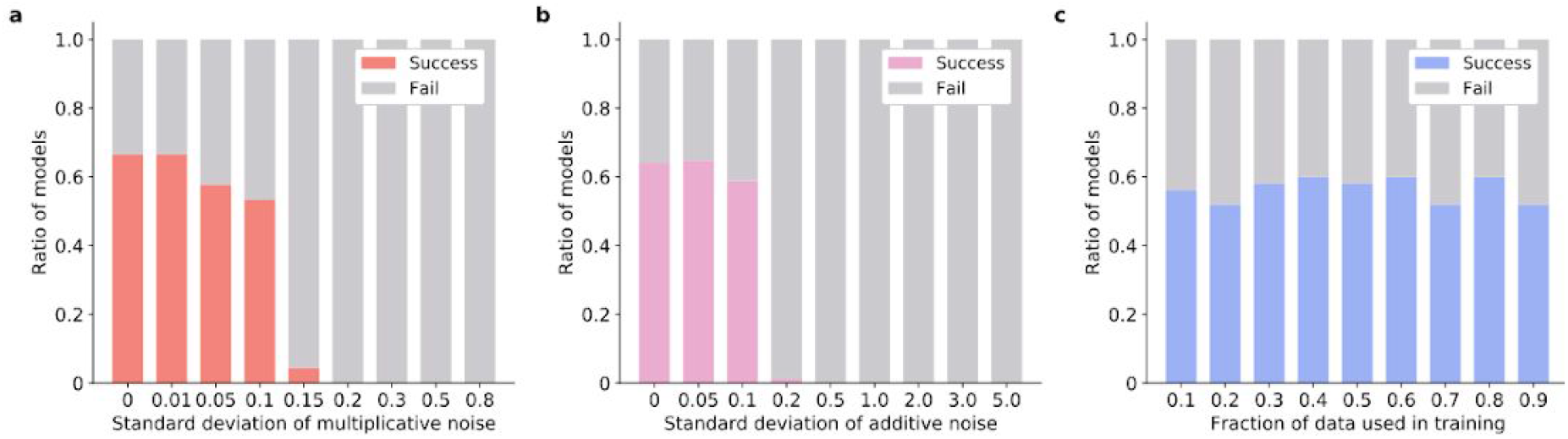
Model convergence against noise and reduced training set. **a-b**. The percentage of models that successfully converged, defined by MSE of training set below a threshold of 0.05, decreases as an increased level of multiplicative Gaussian noise (**a**) or additive Gaussian noise (**b**) was added into training data. Such a decrease in model accuracy is plausible: as the signal is overwhelmed by the increasing noise level, model performance and stability should decrease in terms of MSE. **c.** The percentage of successful models stayed the same as more data was used for model training.

**Figure S9.**
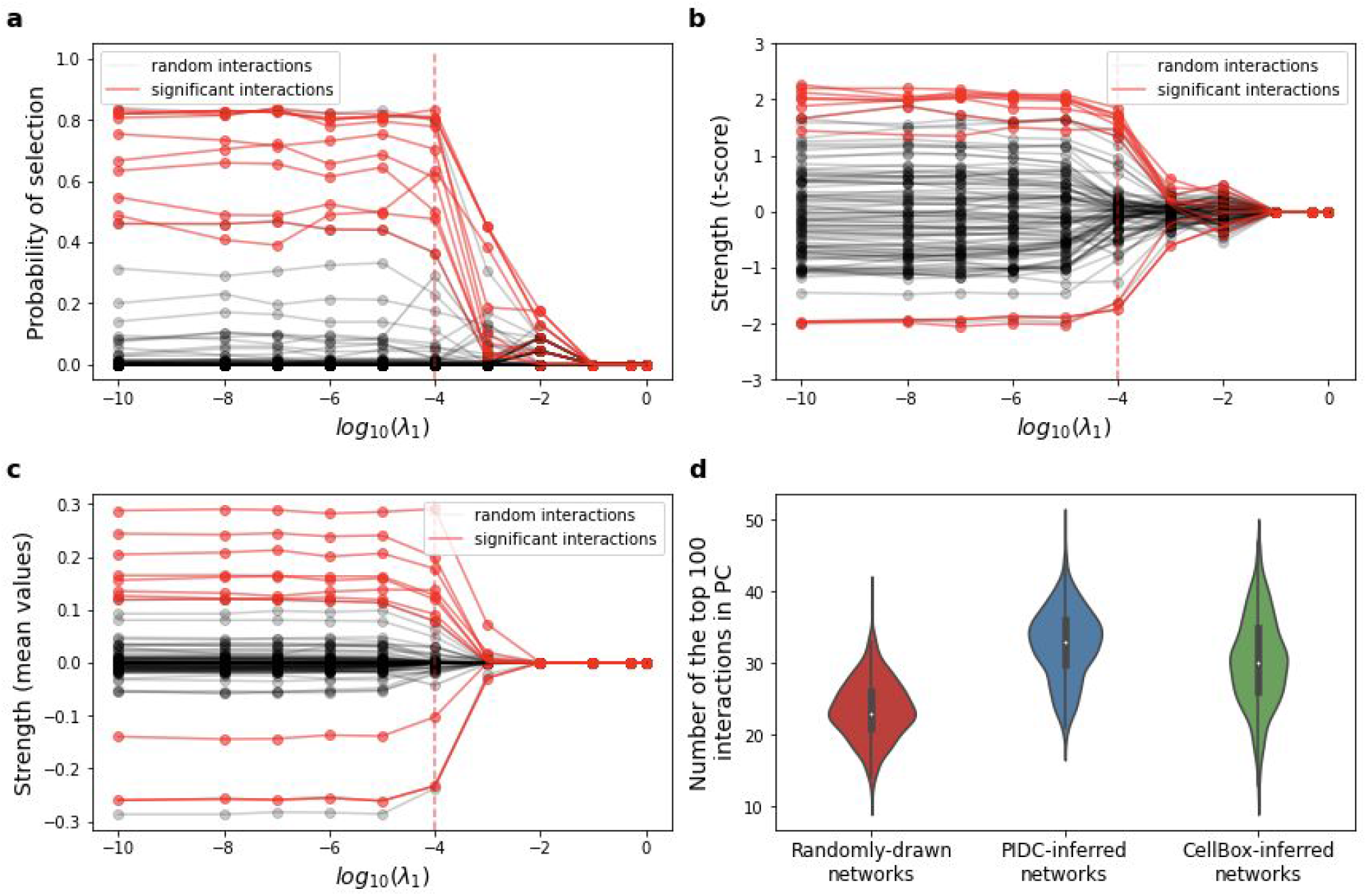
De novo inferred interactions are both robust and significantly consistent with literature derived networks. **a-c**. We used stability selection approaches (Meinshausen and Bühlmann, 2010) to examine network inference stability. We trained CellBox models with different L1 regularization strengths λ_1_ and plotted the stability paths, including the probability of selection (**a**), t-score of the interactions over the models (**b**), and mean values of the interactions over the models (**c**), to compare the top 100 interactions (10 out of these are highlighted in red lines) identified by CellBox to 100 randomly chosen interactions from the pool of all possible (phospho)protein-(phospho)protein interactions (grey lines). The CellBox models result in significantly stable parameter inference, compared to random permutations (m>140 models for each λ_1_). **d.** To quantitatively evaluate our models in the context of prior knowledge, we compare CellBox models to random models: the same number of network models (m = 1,000) were generated with interaction parameters randomly drawn from the pool of all (phospho)protein-(phospho)protein interaction parameters in the CellBox-inferred network models. For each of the inferred or random network models, the interactions existing in PC were identified out of the top one-hundred interactions (ranked by absolute interaction strengths). Our results find a significant difference in the number of interaction edges consistent with prior knowledge from Pathway Commons between the CellBox-inferred network and a network with random interactions (t-test, p=3e-149, N_CellBox-inferred interactions in PC_ - N_random interactions in PC_ = 7). We also compared our methods with other network inference methods (Chan, Stumpf and Babtie, 2017). Partial information decomposition and context (PIDC) recovers more prior knowledge interactions in PC compared to CellBox ^(by t-test, p-value=8.0e-21, N^PIDC-inferred interactions in PC ^- N^CellBox-inferred interactions in PC ^= 2).^ Nevertheless, we argue that the slight advantage of PIDC compared to CellBox in this measure is more than compensated for by the crucial ability of the dynamically executable CellBox model to predict cell response to unseen perturbations, which is the primary objective of our approach.

### Supplementary Tables

Table S1. Annotation of nodes in the network.

Table S2. Model-inferred molecular interactions and their pathway database information

